# A regulatory program for initiation of Wnt signaling during posterior regeneration

**DOI:** 10.1101/2020.03.26.010181

**Authors:** Alyson N. Ramirez, Kaitlyn Loubet-Senear, Mansi Srivastava

## Abstract

Whole-body regeneration requires the re-establishment of body axes for appropriate patterning of new and old tissue. Wnt signaling has been utilized to correctly regenerate tissues along the primary axis in many animals. However, the causal molecular mechanisms that first launch Wnt signaling during regeneration are poorly characterized. We used the acoel worm *Hofstenia miamia* to identify processes that initiate Wnt signaling. Transcriptome profiling, *in situ* hybridization, and functional studies revealed a Wnt ligand, *wnt-3*, as an early wound-induced gene specifically activated in posterior-facing wound sites and was required for establishing posterior identity during regeneration. *wnt-3* was upregulated upon amputation in stem cells, and its inhibition affected stem cell proliferation. Ectopic expression of anterior markers in *wnt-3* RNAi head fragments was stem cell dependent. Chromatin accessibility data revealed that *wnt-3* activation during regeneration required input from the general wound response. Additionally, the expression of a different Wnt ligand, *wnt-1*, prior to amputation was required for activation of wound-induced *wnt-3* expression. Our study establishes a gene regulatory network for initiating Wnt signaling in posterior tissues in a bilaterian.

## Introduction

Animals capable of whole-body regeneration can replace any missing cell type and re-establish entire body axes. Axial repatterning enables regenerated tissues to acquire correct identities according to their locations in the body plan of the animal. In bilaterians, *i*.*e*., animals with distinct anterior-posterior and dorsal-ventral axes, transverse amputation creates two fragments: a “head” fragment with a posterior-facing wound site that must regenerate tail tissue, and a “tail” fragment with an anterior-facing wound site that must regenerate head tissue. Wound sites generated by a single amputation therefore must initiate and establish distinct anterior and posterior regeneration programs, despite having similar positional identities prior to amputation. Mechanistic studies of whole-body regeneration in planarians and acoels, two distantly-related bilaterian species (Figure 1A), identified a requirement for Wnt signaling in posterior regeneration (Gurley, Rink and Sanchez Alvarado, 2008; Iglesias *et al*., 2008; Petersen and Reddien, 2008, 2009; Srivastava *et al*., 2014). Wnt ligands are highly expressed in posterior tissues in the planarian *Schmidtea mediterranea* and in the acoel *Hofstenia miamia* and inhibition of Wnt signaling during regeneration causes the transformation of posterior tissues to anterior structures in both species, giving rise to double-headed animals with head tissue forming at both anterior- and posterior-facing wound sites. Conversely, overactivation of Wnt signaling via inhibition of Wnt antagonists during regeneration gives rise to double-tailed animals in both species. Thus, Wnt signaling represents a putatively conserved mechanism for establishing the identity of posterior tissues in regenerating bilaterians. Furthermore, Wnt signaling is activated specifically at oral-facing wound sites in *Hydra* and is required for correct regeneration of tissues along the oral-aboral axis, the primary body axis in cnidarians(Nakamura *et al*., 2011; Vogg *et al*., 2019). Given the phylogenetic position of cnidarians as the sister-lineage to bilaterians, it is likely that the role of Wnt signaling in patterning the axial identities of tissues during regeneration could be broadly conserved across metazoans (Figure 1A). A robust assessment of this hypothesis of conservation of the role of Wnt signaling during regeneration requires an understanding of how Wnt signaling is initiated upon amputation.

**Figure 1:**
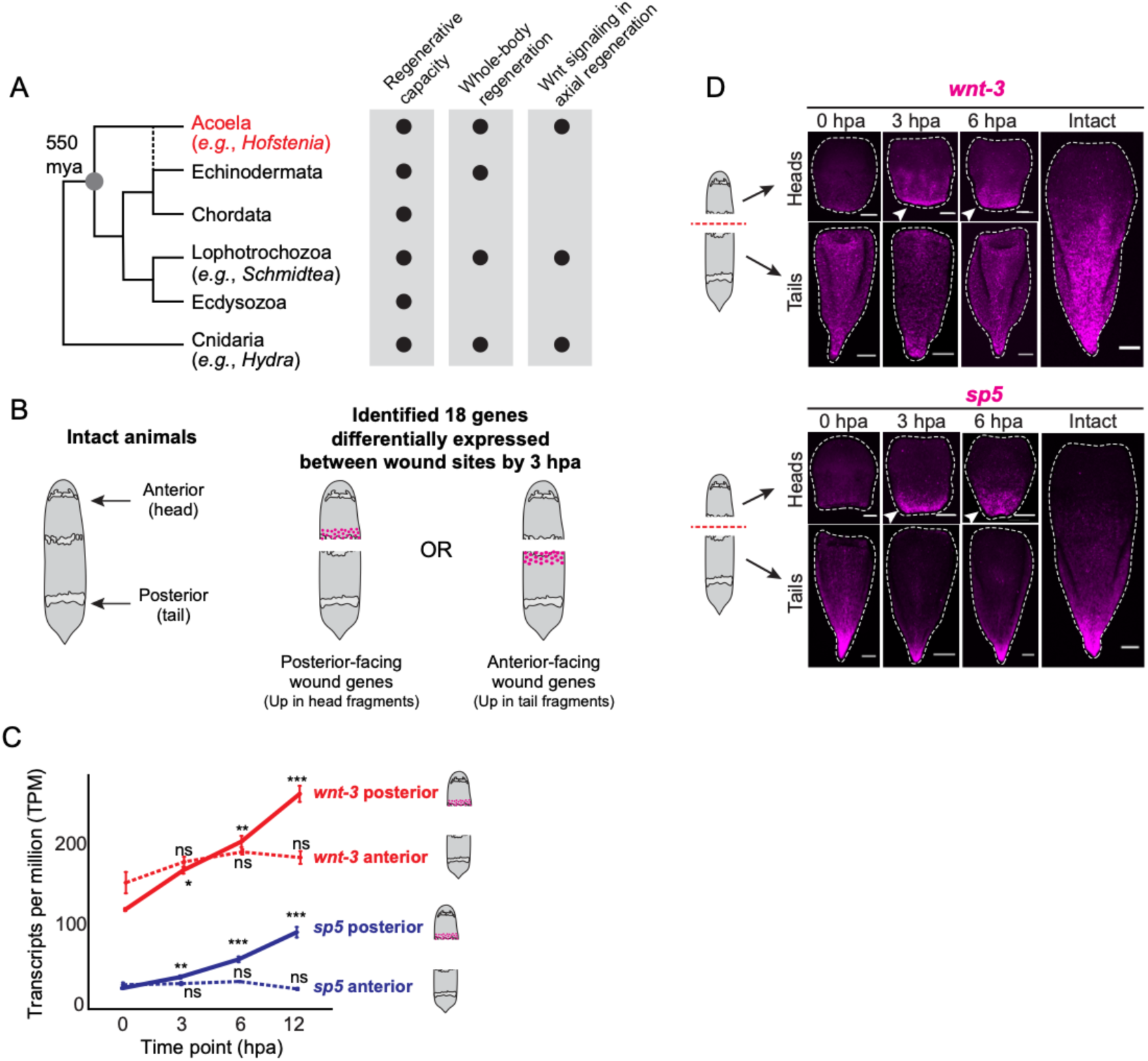
Transcriptomic analysis of regenerating animals identified genes induced asymmetrically at anterior-facing or posterior-facing wound sites. (A) Phylogenetic tree showing the position of acoels such as *Hofstenia* (red text) as a sister-lineage to all other bilaterians, with regenerative capacity and prior knowledge of Wnt utilization in this process. Dashed line indicates a putative relationship between acoels and echinoderms based on an alternative phylogenetic position of acoels as sister to Ambulacraria, proposed by some studies (Philippe *et al*., 2007, 2011). Black circle, trait is present in at least one species of the lineage shown. (B) Schematics of worms showing naming convention used in this paper to denote anterior- or posterior-facing wound sites, and depicting identification of asymmetrically-expressed, early wound-induced genes using a published transcriptome (Gehrke, et al 2019). (C) Transcript per million (TPM) values of *wnt-3* and *sp5* in anterior- and posterior-facing wound sites. (*p*-values, likelihood ratio test; * = *p* < 0.001; ** = p < 0.0001; *** = *p* < 0.00001; ns, not significant). (D) *in situ* hybridization expression patterns for *wnt-3* and *sp5* at various hours post amputation (hpa): 0 hpa (control), 3 hpa, 6 hpa, and intact worms. *wnt-3* and *sp5* were expressed specifically at posterior-facing wound sites during regeneration (arrowheads). Scale bars, 100μm.

In *Hydra*, two distinct phases of Wnt pathway upregulation in oral-facing wound sites are known – immediate secretion of Wnt3 protein by apoptotic interstitial cells and later, transcriptional upregulation of *Wnt3* mRNA observed in endodermal epithelial cells(Hobmayer *et al*., 2000; Guder, 2006; Chera *et al*., 2009; Nakamura *et al*., 2011). Studies of the *Wnt3* locus showed that a combination of activation via β-catenin/TCF and repression via Sp5 restricts the Wnt signaling center to the oral end of intact animals(Nakamura *et al*., 2011; Vogg *et al*., 2019). Although it has been hypothesized that transient suppression of the repressor must occur to enable *Wnt3* activation at oral-facing wound sites(Nakamura *et al*., 2011), and expression analysis shows a corresponding absence of *Sp5* prior to *Wnt3* activation(Vogg *et al*., 2019), the mechanisms leading to transcription of the *Wnt3* locus upon amputation are not known in cnidarians.

It is also unknown which transcriptional programs induce Wnt ligand expression upon wounding in bilaterians. In planarian regeneration, mechanisms for inhibition of Wnt signaling specifically at anterior-facing wound sites have been identified(Gurley, Rink and Sanchez Alvarado, 2008; Iglesias *et al*., 2008; Petersen and Reddien, 2008, 2009; Gaviño *et al*., 2013; Roberts-Galbraith and Newmark, 2013; Tewari *et al*., 2018), but pathways for initiation of Wnt ligand expression, which occurs at both anterior- and posterior-facing wound sites, are yet to be identified. The control of Wnt signaling during regeneration has not yet been investigated in acoels. Here, we sought to assess the dynamics of Wnt pathway expression during regeneration and to identify mechanisms that drive its activation upon amputation in *Hofstenia*.

Our analysis of the regeneration transcriptome of *Hofstenia* revealed that Wnt ligands and other posterior markers are expressed at posterior-facing wound sites within six hours following amputation. To find candidate genes for the initiation of this expression, we focused on the earliest asymmetries between anterior- and posterior-facing wound sites during regeneration. We found that a combination of a generic wound response factor and a pre-existing patterning gradient activates a Wnt ligand specifically at posterior-facing wound sites within three hours upon wounding. Specific establishment of Wnt signaling at posterior-facing wound sites is a shared mechanism for determining correct anterior-posterior specification; our work has identified a regulatory program for the initiation of Wnt signaling during posterior regeneration.

## Results

### Transcriptomic analysis of regenerating animals identified genes induced at either anterior-facing or posterior-facing wound sites

In order to identify regulators for initiating Wnt signaling, which is localized to posterior tissues, we sought to understand the dynamics of symmetry breaking during regeneration. We reanalyzed a transcriptome profiling dataset and noted that known anterior and posterior markers were significantly upregulated by 12 hours post amputation (hpa), with most posterior markers significantly upregulated by 6 hpa (Supplemental Figure 1.1, Gehrke *et al*., 2019). Wound-induced genes identified in *Hofstenia* thus far were found to be upregulated in both anterior- and posterior-facing wound sites. We reasoned that genes involved in initiation of Wnt signaling would be expressed asymmetrically between anterior- and posterior-facing wound sites, and focused on identifying genes in this dataset with asymmetric expression by 3 hpa (Figure 1B). Anterior- and posterior-facing wound sites had ten and eight uniquely-expressed genes respectively, that were upregulated by 3 hpa (Figure 1C, Supplemental Table 1, Supplemental Figure 1.2). These genes included transcription factors known in other systems to regulate posterior identity (both during development and regeneration; *brachyury, sp5*), factors in known signaling pathways (*smoothened, hes, wnt-3*), and other transcription factors (*foxa1*) (Singer *et al*., 1996; Kavka and Green, 1997; Yamaguchi *et al*., 1999; Arnold *et al*., 2000; Estella *et al*., 2003; Thorpe, Weidinger and Moon, 2005; Weidinger *et al*., 2005; Fujimura *et al*., 2007; Martin and Kimelman, 2008; Sun *et al*., 2008; Morley *et al*., 2009; Augello, Hickey and Knudsen, 2011; Srivastava *et al*., 2014; Kennedy *et al*., 2016; Tewari *et al*., 2019).

To validate our transcriptomic analysis, we assessed the expression of all 18 genes with *in situ* hybridization in intact animals and in regenerating fragments at 0 hpa, 3 hpa, and 6 hpa (Figure 1D, Supplemental Figure 1.3). Of the genes that showed visible expression, a quarter (4/16; 25%) did not show wound-induced expression, whereas half (8/16; 50%) were expressed in both anterior- and posterior-facing wound sites. Four genes (*ptn14, brachyury, sp5*, and *wnt-3*) were asymmetrically upregulated between anterior- or posterior-facing wound sites at 6 hpa. Two genes, *sp5* and *wnt-3*, emerged as the only candidates with expression visible by 3 hpa (Figure 1D). Both genes were specifically upregulated at posterior-facing wound sites and are mediators of Wnt signaling(Tewari *et al*., 2019). We therefore focused on studying these two genes to identify mechanisms for Wnt re-establishment in the posterior.

We further validated the dynamics of expression of *sp5* and *wnt-3* during regeneration via quantitative PCR (qPCR) and by extending the time course of expression analysis via *in situ* hybridization (Supplemental Figure 1.4). Both genes were upregulated in posterior-facing wound sites by 3 hpa and maintained expression in the posterior as head fragments continued to regenerate. Further, posterior expression of *sp5* and *wnt-3* was maintained in a gradient in tail fragments, consistent with their known expression in the posterior of intact worms(Srivastava *et al*., 2014; Tewari *et al*., 2019). Therefore, we next asked if *wnt-3* and *sp5* play a role in determining the axial identity of tissue during regeneration in *Hofstenia*.

### wnt-3 RNAi animals failed to regenerate and showed defects in axial polarity

To assess the role of *wnt-3* and *sp5* during regeneration, we used RNA interference (RNAi) to inhibit gene expression prior to transverse amputation and assessed the capacity of RNAi fragments to regenerate by 8 days post amputation (dpa), when head and tail marker expression is re-established. We predicted that if *sp5* or *wnt-3* were required for the re-establishment of axial identity during regeneration, polarity defects would be present following RNAi. *sp5* RNAi fragments inconsistently led to defects with anterior (11/58; 18.9%) and posterior (21/56; 37.5%) regeneration (Supplemental Figure 2.1A). In addition, expression of anterior and posterior markers was not affected following *sp5* knockdown at 8 dpa (Supplemental Figure 2.1B). Notably, RNAi of *wnt-3* led to striking regeneration deficient phenotypes: head fragments failed to form posterior tissues (135/142; 95%), whereas tail fragments failed to regenerate a visible blastema or mouth (136/139; 97.8%) by 8 dpa (Figure 2A, Supplemental Figure 2.2). We therefore focused on further characterizing the *wnt-3* RNAi phenotype.

**Figure 2:**
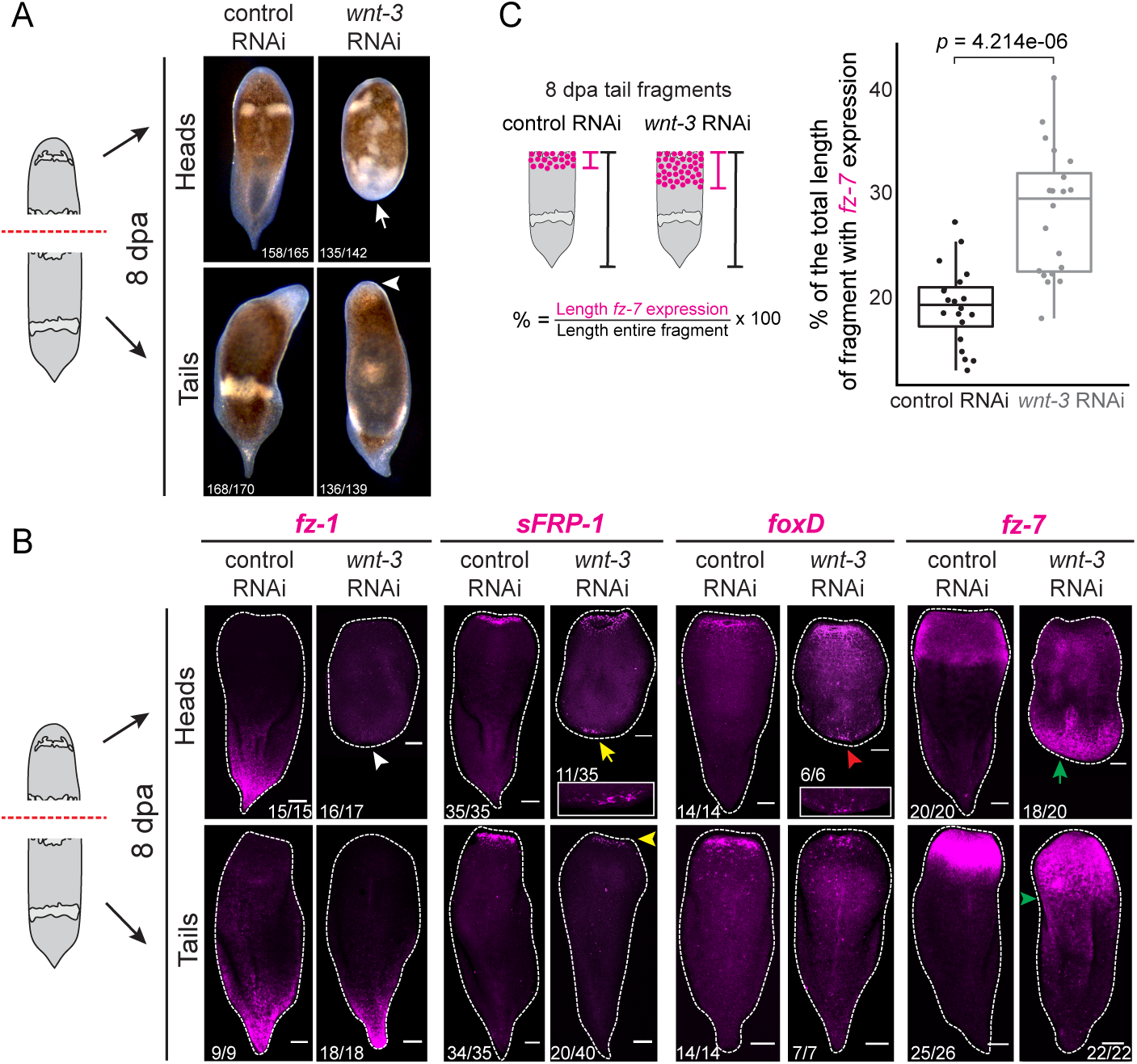
*wnt-3* RNAi animals failed to regenerate and showed defects in axial polarity. (A) Control and *wnt-3* RNAi fragments at 8 days post amputation (dpa). *wnt-3* RNAi head fragments fail to make new posterior structures (arrow), and tail fragments failed to form an unpigmented blastema (arrowhead). (B) Expression of anterior (*sFRP-1, foxD, fz-7*) and posterior (*fz-1*) markers in control and *wnt-3* RNAi head and tail fragments. Expression of *fz-1* was lost in posterior-facing wound sites of *wnt-3* RNAi head fragments (white arrowhead). *sFRP-1* was expressed in posterior-facing wound sites of *wnt-3* RNAi head fragments (yellow arrow, shown magnified in inset) and was either diminished in anterior-facing wound sites of tail fragments (20/40; yellow arrowhead) or completely lost (20/40; data not shown). *foxD* expression was detected at posterior-facing wound sites of *wnt-3* RNAi head fragments (red arrowhead, shown magnified in inset). *fz-7* was expressed in posterior-facing wound sites of *wnt-3* RNAi head fragments (green arrow). In *wnt-3* RNAi tail fragments, *fz-7* was expanded towards the posterior (green arrowhead). Proportions of animals with phenotype imaged in the lower left corner. Scale bars, 100μm. (C) Quantification of the expansion of *fz-7* in *wnt-3* RNAi tail fragments compared to control. (n = 20 fragments/RNAi condition; *p*-value < 0.00001, Welch two-sample *t*-test).

To determine if attenuation of *wnt-3* during regeneration resulted in defects in polarity, we assessed the expression of anterior and posterior markers in *wnt-3* RNAi fragments. *wnt-3* RNAi head fragments expressed anterior markers (*sFRP-1, foxD*, and *fz-7*) within posterior-facing wound sites, and failed to express the posterior marker *fz-1*, indicating a mis-specification of the posterior-facing wound site (Figure 2B). *wnt-3* RNAi tail fragments showed some expression of the anterior markers *sFRP-1* and *foxD* at the anterior-facing wound site, but relative to control RNAi, the level of expression appeared reduced (Figure 2B). The expression domain of *fz-7* at anterior-facing wound sites in *wnt-3* RNAi was expanded compared to control RNAi tails, indicating that *wnt-3* is required for restricting expression of anterior markers (Figure 2C). Taken together, the *wnt-3* RNAi phenotype suggests RNAi fragments both (1) failed to correctly specify a new posterior and (2) correctly specified the anterior, but failed to achieve full anterior regeneration by 8 dpa.

The misexpression of anterior markers at posterior-facing wound sites following *wnt-3* RNAi resembled the RNAi phenotype of a previously-studied Wnt ligand, *wnt-*1 (Srivastava et al., 2014). However, there are notable differences between regenerating *wnt-3* and *wnt-1* RNAi animals. First, *wnt-1* RNAi tail fragments made a full blastema and a visible mouth by 8 dpa (Supplemental Figure 2.3A), which *wnt-3* RNAi tail fragments failed to achieve (Figure 2A). Second, although both *wnt-3* and *wnt-1* RNAi head fragments expressed anterior markers ectopically within posterior-facing wound sites, *wnt-1* fragments formed a complete ectopic head with a clear mouth, as assessed by expression of *sFRP-1* (Supplemental Figure 2.3A). In contrast, *wnt-3* RNAi head fragments expressed *sFRP-1* in the posterior-facing wound sites, but these cells did not form coherent structures (Figure 2B). Third, in anterior-facing wound sites, *wnt-1* RNAi tail fragments had no discernible expansion of the *fz-7* expression domain (Supplemental Figure 2.3A). Conversely, following *wnt-3* RNAi, expression of *fz-7* is expanded in anterior-facing wound sites (Figure 2B, C). Fourth, the expression of *wnt-1* and *wnt-3* during regeneration is distinct. Although *wnt-1* did not emerge as an early posterior wound-induced gene in our transcriptome analysis, it did show an upward trend in expression in posterior wound sites (Supplemental Figure 2.3B). However, this was not validated in our experimental studies as we did not detect wound-induced expression of *wnt-1* during regeneration by *in situ* hybridization or by qPCR (Supplemental Figure 2.3C, D). In contrast, the earliest expression of *wnt-3* in posterior-facing wound sites is wound-induced, and is detectable by 3 hpa in all three measurements (Figure 1C, D, Supplemental Figure 1.4C, D). Taken together, these results suggest distinct roles for *wnt-3* and *wnt-1* during regeneration.

### wnt-3 expression and function involves stem cells

RNAi animals failed to make both anterior and posterior outgrowths. We sought to determine if this phenotype was the result of a general defect in cell proliferation in *wnt-3* RNAi animals, and if a lack of proliferation could underlie the defects we found following *wnt-3* RNAi. In *Hofstenia*, similar to planarians, the only proliferative cells are the neoblasts, a population of effectively pluripotent stem cells that are required for regeneration(Reddien *et al*., 2005; Srivastava *et al*., 2014). Phospho-histone H3 immunostaining previously revealed dynamic changes in neoblast proliferation during regeneration in *Hofstenia*, therefore we utilized this assay to assess the number of mitotic cells in *wnt-3* RNAi fragments (n > 4 fragments/time point). We detected a decrease in the number of proliferating cells in both head and tail fragments specifically close to the wound site at 72 hpa (Figure 3A, Supplemental Figure 3.1). This indicates the expression of *wnt-3* is required for cell proliferation within regenerating fragments, and implies a wider role for Wnt signaling in regulating stem cell proliferation in response to amputation. However, the inability to correctly pattern posterior-facing wound sites in *wnt-3* RNAi cannot be explained by a complete loss of cell proliferation at this wound site.

**Figure 3:**
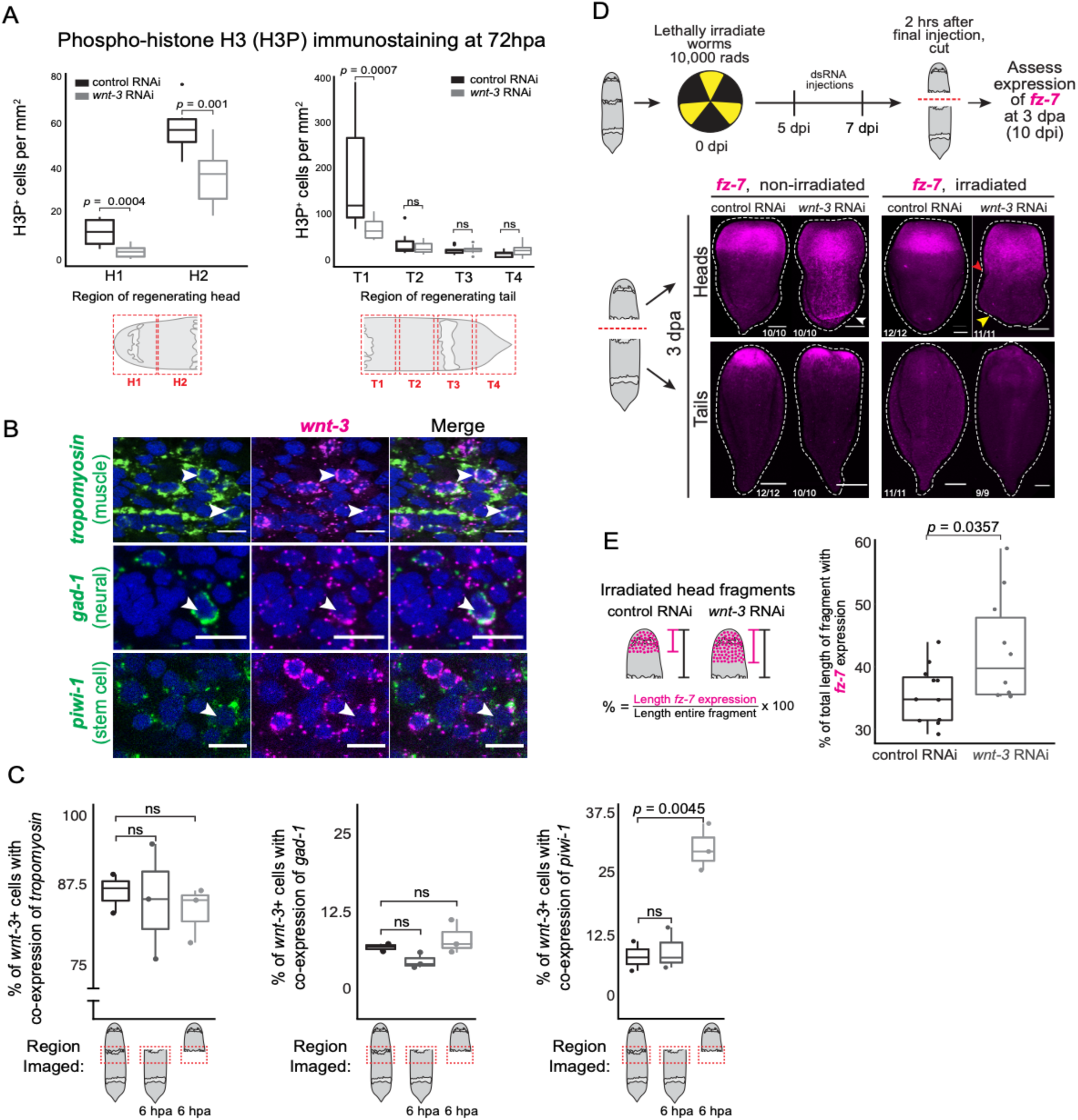
wnt-3 expression and function involves stem cells. (A) Quantification of H3P+ foci in head and tail fragments rol and wnt-3 RNAi animals at 72 hpa. Because H3P+ foci showed regionalized distribution, H3P+ foci per unit area ompared by region (head fragments, 2 regions; tail fragments, 4 regions; n = 10 fragments/RNAi condition; p-value, on rank-sum test; ns, not significant). (B) Representative images of cell-type expression patterns for wnt-3, assessed expression with tissue markers (tropomyosin, muscle; piwi-1, stem cells; gad-1, neural) in intact worms or 6 hpa rating fragments. White arrows, examples of co-expressed cells. Scale bar, 10μm. (C) Quantification of cell co-sion at either anterior- or posterior-facing wound sites, and an equivalent site in whole animals (p-value, Shapiro-Wilks ity test and Welch two-sample t-test; ns, not significant). (D) Schematic of irradiation experiment. Animals were first irradiated, then at five days post irradiation (dpi) were injected with dsRNA for three consecutive days, amputated, en assessed for expression of fz-7 at 3 dpa/10 dpi. wnt-3 RNAi head fragments expressed fz-7 within posterior-facing sites (white arrowhead). Expression of fz-7 at posterior-facing wound sites in wnt-3 RNAi heads is lost following ion (yellow arrowhead), but the pre-existing domain of fz-7 expression showed expansion toward the posterior (red ead). Scale bar, 100μm. (E) Quantification of pre-existing fz-7 expression in irradiated head fragments (n > 10 nts/condition; p-value, Welch two-sample t-test).

After finding a striking reduction in cell proliferation following *wnt-3* RNAi, we sought to determine a mechanism for *wnt-3* action during regeneration by asking which tissue types expressed *wnt-3* in intact and regenerating animals. To assess if *wnt-3* was expressed in stem cells, we quantified the co-expression of *wnt-3* with a known neoblast marker (*piwi-1*). Because *wnt-3* is known to be expressed in *Hofstenia* muscle(Raz *et al*., 2017), we also assessed co-expression of *wnt-3* and a muscle marker (*tropomyosin*), and a tissue with no previously known Wnt expression (neural; *gad-1*) in intact and 6 hpa regenerating head and tail fragments by *in situ* hybridization. *wnt-3* was expressed in all three cell populations we assessed, in both intact and regenerating animals (Figure 3B, Supplemental Figure 3.2). Notably, significantly more *wnt-3*^*+*^ cells also expressed *piwi-1* in posterior-facing wound sites at 6 hpa compared to a similar region within intact animals (Welch two-sample *t*-test; *p*-value, 0.0045; Figure 3C; Supplemental Figure 3.2; Supplemental Table 1). As *wnt-3* is highly upregulated at 6 hpa within posterior-facing wound sites, the increased proportion of *wnt-3*^*+*^*/piwi-1*^+^ cells suggests *wnt-3* is wound-induced in the *piwi-1*^+^ population.

Given *wnt-3* is expressed within *piwi-1*^+^ stem cells and *wnt-3* RNAi reduced stem cell proliferation, we next asked if the misexpression of anterior markers following *wnt-3* RNAi, *i*.*e*., ectopic *fz-7* expression in posterior-facing wound sites, relied upon stem cells. We irradiated intact *Hofstenia* to ablate the stem cell population and assessed the expression of *fz-7* within regenerating *wnt-3* RNAi fragments. An effect on expression of *fz-7* in the posterior-facing wound site of *wnt-3* RNAi animals would imply that new cell formation or signaling from stem cells within the regenerating fragment contribute to this phenotype. Irradiated worms were injected with dsRNA to inhibit *wnt-3* at 5 days post irradiation (dpi) for three consecutive days and were assessed for expression of the anterior marker *fz-7* at 3 dpa/10dpi (Figure 3D). The posterior-facing wound site expression of *fz-7* in *wnt-3* RNAi heads was lost following irradiation, indicating that stem cells contribute to the *wnt-3* RNAi phenotype (Figure 3D, Supplemental Figure 3.3) either through their progeny or via an unknown signaling mechanism. Interestingly, the anterior expression of *fz-7* within head fragments was expanded following *wnt-3* RNAi (Figure 3E), implying *fz-7* expression can also be reshaped in head fragments following *wnt-3* RNAi without stem cell contribution. Taken together, these observations suggest both stem cells and pre-existing cells contribute to the ectopic expression pattern of the anterior marker *fz-7* within *wnt-3* RNAi fragments.

### Input from the generic wound response is required for wnt-3 expression at posterior-facing wound sites

Once we established that *wnt-3* is expressed asymmetrically soon after amputation and that it is required for correct establishment of posterior identity during regeneration, we sought to determine the mechanism for *wnt-3* activation during regeneration. To identify candidate regulators of *wnt-3* expression, we reanalyzed published ATAC-seq data from regenerating fragments (Figure 4A; Gehrke *et al*., 2019). Using this dataset, we examined the *wnt-3* locus for differentially accessible regions within the promoter and +/- 5kb (10kb total) around the gene locus and studied binding sites for known transcription factors contained within these regions. In both anterior- and posterior-facing wound site datasets, we found a region in the promoter of the *wnt-3* locus that shows a significant increase (adjusted *p*-value, 3.1×10^−8^, tail dataset; Wald test) in accessibility at 6 hpa compared to 0 hpa. We noted the presence of two binding sites for the general wound response factor EGR within this region. Accessibility of this region is significantly reduced (adjusted *p*-value, 0.005, Wald test) following *egr* RNAi relative to control RNAi, implying *wnt-3* is transcriptionally regulated by Egr during regeneration (Figure 4B).

**Figure 4:**
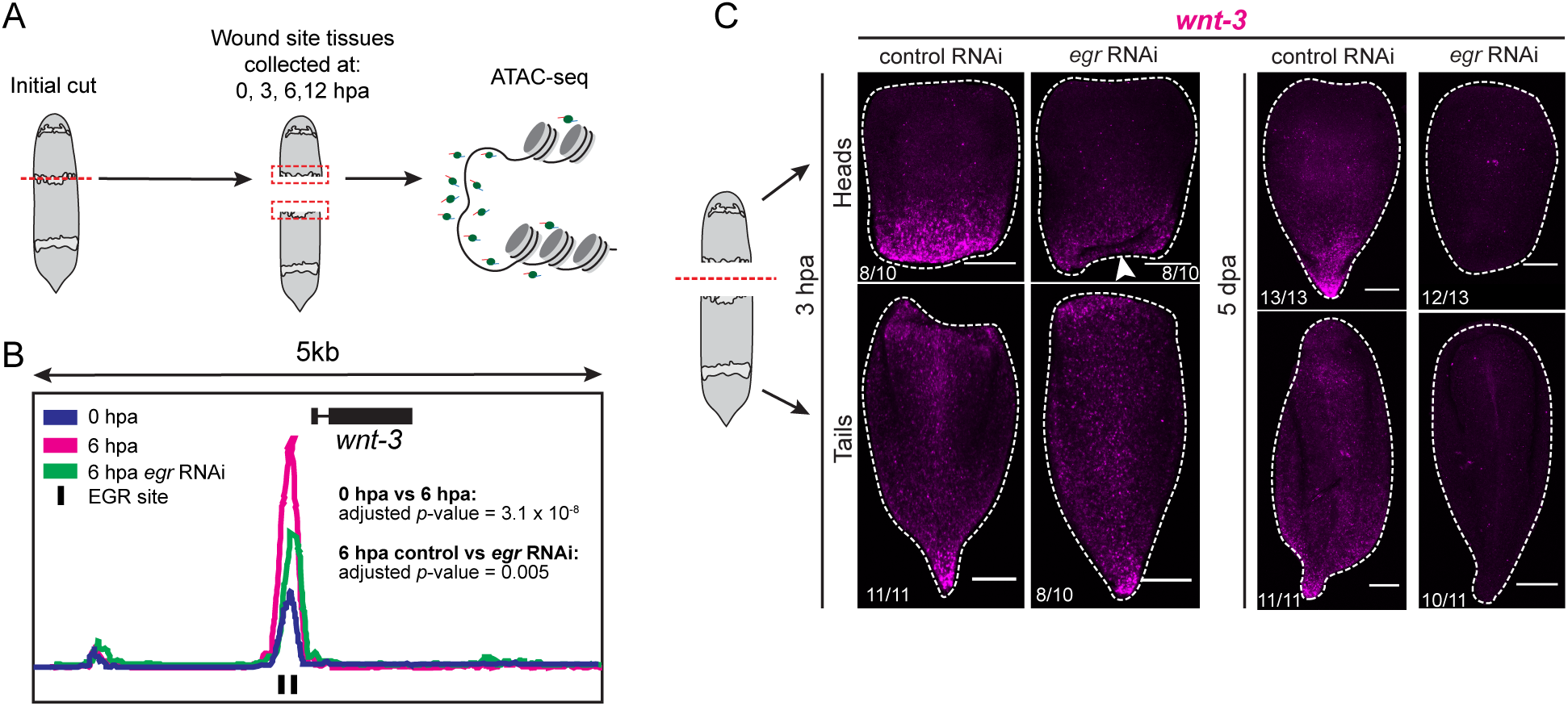
Input from the generic wound response is required for *wnt-3* expression at posterior-facing wound sites. A) Schematic of ATAC-seq data collection (Gehrke et al, 2019). (B) Schematic of the *wnt-3* genomic locus with ATAC-seq data mapped. The promoter region contained a regeneration-responsive peak that is variable from 0 hpa (blue) to 6 hpa (magenta) (adjusted *p*-value, 3.1×10^−8^, tail dataset; Wald test). This peak contained two EGR binding sites (black lines) and the amplitude of this peak was significantly reduced in *egr* RNAi (green) relative to control RNAi (not shown) (adjusted *p*-value, 0.005, Wald test). Control RNAi track has been omitted for clarity. (C) RNAi of *egr* led to loss of wound-induced *wnt-3* expression in regenerating heads (arrowhead) at 3 hpa. By 5 dpa, *wnt-3* expression is lost from both regenerating head and tail fragments in *egr* RNAi animals compared to control RNAi animals. Scale bars, 100μm

To test the hypothesis that *egr* is required for wound-induced expression of *wnt-3*, we performed *egr* RNAi and assessed expression of *wnt-3* during regeneration. The wound-induced expression of *wnt-3* at posterior-facing wound sites was diminished relative to controls by 3 hpa, and was completely lost by 5 dpa (Figure 4C). Notably, pre-existing *wnt-3* expression in tail fragments was also lost by 5 dpa. This suggests that *wnt-3* is transcriptionally regulated by an early wound response factor both for wound-induced expression and for maintenance of expression during regeneration. Furthermore, the locus of *wnt-1*, the other Wnt ligand with a known role in mediating regeneration polarity, did not show a dynamic region of chromatin under the control of *egr*, which is consistent with the absence of detectable wound-induced expression of *wnt-1* at early time points in regeneration (Supplemental Figure 2.3E). We therefore propose that *wnt-3*, and not *wnt-1*, relays the decision-making that enables *Hofstenia* to begin regenerating posterior tissues at posterior-facing wound sites.

### Pre-existing patterning information is required for wnt-3 expression at posterior-facing wound sites

Because *egr* is upregulated at both anterior- and posterior-facing wound sites(Gehrke *et al*., 2019), the control of *wnt-3* by *egr* does not explain the asymmetric, early wound-induced expression of *wnt-3*. Other factors must feed into this locus either to specifically upregulate *wnt-3* at posterior-facing wound sites, or to downregulate *wnt-3* at anterior-facing wound sites. To assess a potential source of information that results in asymmetric *wnt-3* expression as early as 3 hpa, we asked if pre-existing gradients of Wnt pathway members with known effects on regeneration polarity could play a role in establishing this asymmetry.

First, we asked if *wnt-1* was required for asymmetric upregulation of *wnt-3* expression in posterior-facing wound sites. To assess this, we profiled *wnt-3* expression during regeneration using *in situ* hybridization in *wnt-1* RNAi fragments. Following *wnt-1* RNAi, we did not observe wound-induced *wnt-3* expression at posterior-facing wound sites (Figure 5A, Supplemental Figure 5.1A). Although *wnt-1* is expressed in amputated head fragments, this expression corresponds to the levels of *wnt-1* mRNA present in this region of the worm prior to amputation. Because *wnt-1* was not detected as wound-induced by 3 hpa via *in situ* hybridization, transcriptome profiling, or qPCR, we reasoned that the pre-existing expression of *wnt-1* is responsible for activation of *wnt-3* specifically in posterior-facing wound sites at 3 hpa (Supplemental Figure 2.3B, C, D).

**Figure 5:**
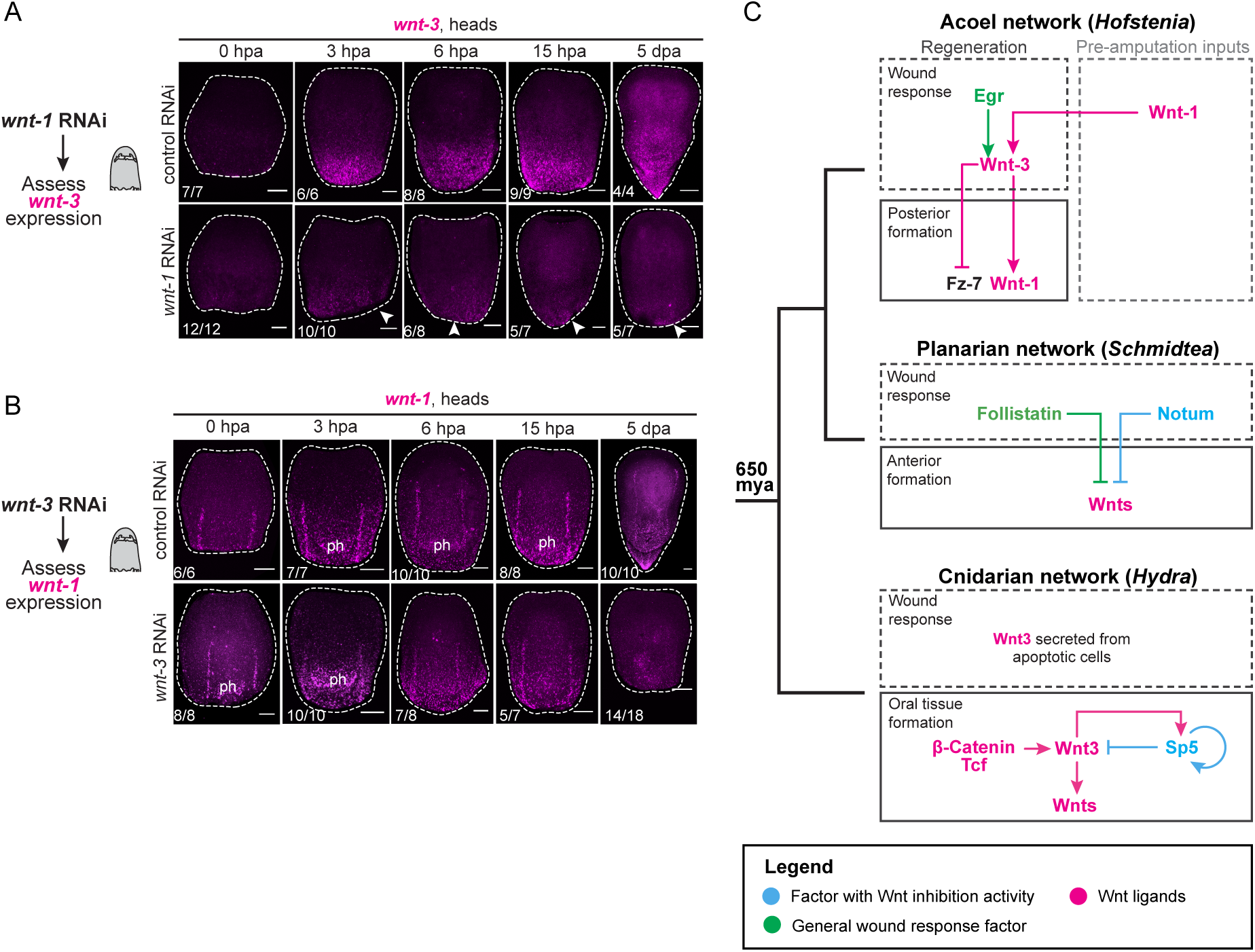
Pre-existing patterning information is required for *wnt-3* expression at posterior-facing wound sites. (A) Expression of *wnt-3* in control and *wnt-1* RNAi regenerating heads. Following *wnt-1* RNAi, *wnt-3* expression was diminished by 3 hpa and failed to be upregulated at the posterior-facing wound site (arrowheads). (B) Expression of *wnt-1* in control and *wnt-3* RNAi heads. RNAi of *wnt-3* did not affect expression of *wnt-1* during the early stages of regeneration, prior to 5 dpa. ph, constitutive expression in pharynx. Scale bars, 100μm. (C) Models for regulation of Wnt signaling upon amputation in acoels (*Hofstenia*), planarians (*Schmidtea*), and cnidarians (*Hydra*). *Hofstenia* and *Hydra* models depict mechanisms for initiation of Wnt ligand expression in posterior and oral (head) tissues respectively; planarian model depicts mechanisms for Wnt inhibition during anterior regeneration. Legend at bottom; phylogenetic tree at left depicts evolutionary relationships between the three species. In *Hofstenia*, Wnt ligands are expressed in posterior gradients prior to amputation. Following amputation, Wnt-3 is upregulated in posterior-facing tissues in response to wounding by the transcription factor Egr; wound-induced expression of Wnt-3 also relies on pre-existing expression of Wnt-1. Wound-induced Wnt-3 expression in turn induces expression of other Wnt ligands (Wnt-1), and represses expression of anterior genes (Fz-7) to promote posterior formation. In planarians, the wound-induced factor Follistatin and the Wnt antagonist Notum work to clear Wnts from anterior-facing wound sites, promoting anterior formation. The inputs activating Wnts in planarians are unknown. In *Hydra*, post-translational regulation of Wnt3 protein leads to its secretion by apoptotic cells specifically in oral-facing wound sites immediately after wounding. Later, *Wnt3* mRNA is transcribed in a β-catenin/TCF-dependent manner and regulated by the head inhibitor Sp5.

In contrast, *wnt-3* RNAi did not affect *wnt-1* expression from 0 hpa to 15 hpa (Figure 5B, Supplemental Figure 5.1B). Yet, *wnt-1* expression in posterior-facing wound sites was lost by 5 dpa following *wnt-3* RNAi, suggesting that *wnt-3* is required for the maintenance of *wnt-1* at later stages of regeneration (Figure 5B). We hypothesize that *wnt-1* and *wnt-3* regulate each other’s expression, albeit in different processes: *wnt-1* is needed for activation of *wnt-3* in posterior-facing wound sites and *wnt-3* is needed for subsequent maintenance of *wnt-1* in regenerating posterior tissue.

Second, we asked if other Wnt pathway components might act to repress *wnt-3* expression at anterior-facing wound sites. In planarians, *notum* is required for setting up correct anterior-posterior identity by clearing Wnt expression from anterior-facing wound sites (Petersen and Reddien, 2011; Roberts-Galbraith and Newmark, 2013). While *notum* expression in *Hofstenia* does not appear to be wound-induced or even expressed in the anterior in intact worms, *notum* RNAi results in two-tailed animals (Supplemental 5.2A,B; Srivastava *et al*, 2014). Thus, we assessed whether the pre-existing expression of *notum* prior to amputation has a role in suppressing *wnt-3* expression at anterior-facing wound sites. Control and *notum* RNAi animals were indistinguishable with regards to *wnt-3* expression at early time points upon amputation in both head and tail fragments (Supplemental Figure 5.2C, D). However, *notum* RNAi tails had robust *wnt-3* expression in anterior-facing wound sites with distinct ectopic tail morphology 5 dpa. This was in contrast with control RNAi tail fragments, which had no *wnt-3* expression at the anterior-facing wound site 5 dpa (Supplemental Figure 5.2C). We propose that whereas *notum* does not regulate wound-induced *wnt-3* expression, it could act more generally by dampening Wnt signals within both head and tail regenerating fragments. While our work has established that *notum* is not required for clearing wound-induced *wnt-3* expression from anterior-facing wound sites, it is plausible that other Wnt regulators or inhibitors could be involved in this process.

Our results suggest a model in which a general wound response factor and pre-existing expression of a Wnt ligand are required for the establishment of asymmetric, wound-induced *wnt-3* expression in posterior-facing wound sites by 3 hpa (Figure 5C). At later time points during regeneration, after the posterior identity decision has been made, *wnt-3* maintains expression of posterior regulators in posterior-facing wound sites in a stem cell dependent manner and restricts the expression of anterior genes.

## Discussion

Our investigation of the mechanism for Wnt pathway activation in the posterior in *Hofstenia* revealed the Wnt ligand *wnt-3* was specifically upregulated at posterior-facing wound sites by 3 hpa and was required for correct patterning of posterior tissues. In addition to showing that *wnt-3* function is stem cell mediated, we inferred a gene regulatory network for the activation of this Wnt ligand in posterior-facing wound sites. Notably, we demonstrate a link between the general early wound response and the initiation of patterning information, showing *wnt-3* is transcriptionally regulated by Egr, an early wound response factor. Given that *egr* is upregulated at both anterior- and posterior-facing wound sites, another input into the *wnt-3* locus is needed to cause the significant expression differences between anterior- and posterior-facing wound sites by 3 hpa. We propose that this input is likely controlled by the pre-existing gradient of another Wnt ligand, *wnt-1*, prior to amputation. The role for *wnt-3* is distinct from that of *wnt-1*, which is needed for posterior regeneration but is not the initiator of the posterior regeneration program.

Wnt signaling centers are re-established in a polarized manner along the primary axis during whole-body regeneration in *Hydra* (cnidarian), *Schmidtea* (planarian), and *Hofstenia* (acoel) (Gurley, Rink and Sanchez Alvarado, 2008; Petersen and Reddien, 2008, 2009; Chera *et al*., 2009; Nakamura *et al*., 2011; Srivastava *et al*., 2014; Gufler *et al*., 2018; Vogg *et al*., 2019). Our study of Wnt pathway re-establishment in *Hofstenia* enables cross-species comparisons of this process (Figure 5C).

First, perturbation of wound-induced Wnt ligands affects cell proliferation in both *Hofstenia* and *Hydra(Chera et al., 2009*). Additionally, the control of *wnt-3* in *Hofstenia* by another Wnt ligand (*wnt-1*) bears similarity to the known regulation of the *Hydra Wnt3* locus via *β-catenin/TCF(Gufler et al., 2018)*. However, the identity of the Wnt ligand mediating control via *β-catenin/TCF* is unknown in *Hydra*.

Second, the direct linkage between a wound response factor and a Wnt locus observed in the *Hofstenia* network (Figure 5) has been hypothesized but not demonstrated in *Hydra*. Control of the *Wnt3* locus in *Hydra* could be through the wound-induced CREB transcription factor, as binding sites are present in this region(Galliot *et al*., 1995; Kaloulis *et al*., 2004; Chera, Kaloulis and Galliot, 2007; Nakamura *et al*., 2011). Based on the phylogenetic positions of cnidarians and acoels, we propose that upregulation of Wnt ligands during regeneration via the combined effect of general wound response factors, Wnt signaling, and the downstream control of cell proliferation could be a general feature of axial regeneration, although the identities of the regulatory factors may differ across species. This hypothesis can be tested by mechanistic studies of regeneration in other bilaterians such as planarians.

Third, the spatial dynamics of Wnt ligand upregulation found in *Hofstenia* are similar to those in *Hydra*, but differ from those in planarians. Early Wnt ligand upregulation upon bisection is identified as asymmetric between the two wound sites in both *Hofstenia* and *Hydra*; in *Hofstenia, wnt-3* is only upregulated in posterior-facing wound sites, and in *Hydra, Wnt3* is only detected at oral-facing wound sites(Lengfeld *et al*., 2009; Nakamura *et al*., 2011). In contrast, in planarians the *wnt1* ligand is upregulated at both anterior- and posterior-facing wound sites(Petersen and Reddien, 2009). Furthermore, in planarians the general wound response factor *follistatin* and the anterior-expressed Wnt antagonist *notum* inhibit Wnt signaling, ultimately restricting Wnt signaling to posterior-facing wound sites(Gaviño *et al*., 2013; Roberts-Galbraith and Newmark, 2013). Further studies of transcriptional control of Wnt ligand re-expression in planarians and of Wnt signaling inhibition in cnidarians (in aboral-facing wound sites) and acoels (in anterior-facing wound sites) are needed for more systematic comparisons across the three species.

Fourth, the utilization of Wnts during regeneration in planarians, acoels, and cnidarians possibly occurs in different cell types. Wnt expression has been previously characterized within muscle tissue in acoels and planarians(Witchley *et al*., 2013; Raz *et al*., 2017; Scimone, Cote and Reddien, 2017). In planarians, expression of Wnt ligands required for posterior identity (*Smed-wnt1, Smed-wntP-2*) are highly enriched in muscle tissue(Witchley *et al*., 2013). In cnidarians, Wnt ligands are expressed within both endodermal and ectodermal epithelial cells and Wnt3 protein is activated and secreted by interstitial stem cells upon amputation(Chera *et al*., 2009; Lengfeld *et al*., 2009; Nakamura *et al*., 2011; Petersen *et al*., 2015; Siebert *et al*., 2019). However, it is unknown if Wnt signaling from these different cell types serves similar or distinct functions. Here, we demonstrated that *Hofstenia* expresses *wnt-3* within stem, neural, and muscle cells, and that the localization of *wnt-3* is enriched in stem cells during regeneration. Further characterization will be necessary to disentangle the roles of *wnt-3* within each tissue.

Wnt signaling has a well-known role in controlling the primary axis in cnidarians (oral-aboral) and bilaterians (anterior-posterior) during development(Hobmayer *et al*., 2000; Martin and Kimelman, 2009; Loh, van Amerongen and Nusse, 2016). The shared utilization of the Wnt pathway for axial regeneration, albeit with potentially different initiation programs or cellular mechanisms during regeneration in cnidarians, planarians, and acoels, which diverged 650 million years ago, could reflect two alternative evolutionary histories. First, the use of Wnts in patterning the body axis could have been independently co-opted in each species, potentially redeployed from their role in axial polarity establishment during development. Second, the differences in the mechanism initiating axial polarity could be indicative of developmental systems drift. In this scenario, the roles of Wnts in patterning body axes during regeneration could have been shared by the last common ancestor, and over time the mechanism to initiate Wnt activity upon amputation could have diverged. Further characterization of the development of each species, the role Wnt signaling plays in this process, and the mechanisms used to re-establish polarity in other animals capable of whole-body regeneration will be required to distinguish between these possibilities.

## Acknowledgements

We thank Dr. Andrew R. Gehrke for insights and help in analyzing chromatin accessibility data and Dr. D. Marcela Bolaños for blind-scoring of experiments. M.S. is supported by the Searle Scholars Program, Smith Family Foundation, and the National Science Foundation (Award #1652104). A.N.R. is supported by the HHMI Gilliam Fellowship.

## Author Contributions

A.N.R. and M.S. designed the study and wrote the manuscript. K.L-S. performed and analyzed cellular co-expression experiments. A.N.R. conducted all other experiments.

## Materials and Methods

### Analysis of RNA-seq data

Dataset was collected as described in(Gehrke *et al*., 2019). Briefly, animals were bisected by cutting at the middle stripe and allowed to regenerate for 0, 3, 6, 12, and 48 hours post amputation (hpa) prior to collection of tissue from the wound site. Anterior- and posterior-facing wound sites were collected separately prior to sequencing. All genes included in this analysis had a minimum transcript per million (TPM) value of 5 at 0 hpa, as values below this could confound the analysis. Genes were identified as wound induced (adjusted *p*-value, < 0.05 at 6 hpa compared to 0 hpa; likelihood ratio test) and filtered for significant expression at 3 hpa compared to 0 hpa (*p*-value < 0.001; likelihood ratio test) within the corresponding head or tail dataset (Supplemental Table 1). We first applied the same adjusted *p*-value thresholds to the head and tail datasets, but this criterion effectively removed all tail genes from our analysis. To ensure that tail genes were included, we loosened the significance thresholds for the tail dataset at 6 hpa (adjusted *p*-value, < 0.5) and 3 hpa (*p*-value < 0.005). Due to the looser criteria for the tail dataset, some genes that were significantly upregulated in the head dataset appeared in the tail list, which was reflected in their expression in our final analysis (Supplemental Figure 1.2) and in their *in situ* hybridization patterns (Supplemental Figure 1.3) at 6 hpa.

### Fluorescent in situ hybridization (FISH)

Whole animals or regenerating fragments were fixed rocking at room temperature for 1 hour in 4% paraformaldehyde solution in PBS+0.1% Triton-X-100. Animals were dehydrated gradually into 100% MeOH and stored at -20°C. FISH protocol was the same as described previously(Srivastava *et al*., 2014; Gehrke *et al*., 2019) with two modifications; (1) all wash steps were completed with 800μL of solution, and (2) animals were bleached for 2hrs at room temperature in bleach solution (5% deionized formamide,1.2% H_2_O_2_, 50% saline-sodium citrate buffer in milli-Q water) under a strong light source. Samples were imaged using a Leica SP8 confocal microscope.

### Quantitative PCR (qPCR)

Wound sites from amputated fragments were collected (replicates = 3, n = 4 fragments/replicate) at multiple time points during regeneration (0, 3, 6, 9, 15, 24, 48 hpa). RNA was extracted using the Nucleospin RNA XS kit (740902.10, Macherey-Nagel). Extracted RNA was used for cDNA synthesis using the SuperScript III kit (Thermo Fisher <u>18080044</u>) with random hexamer priming. Expression of target genes was detected using forward and reverse primers (Supplemental Table 1) and normalized by expression of a housekeeping gene (*gapDH* or *ef1-alpha*). For validation of gene knockdown using qPCR, whole regenerating head or tail fragments were collected at 8dpa (replicates = 3, n = 3 fragments/replicate).

### RNA interference (RNAi)

RNAi was performed as previously described in (Srivastava *et al*., 2014; Gehrke *et al*., 2019). Animals were soaked in dsRNA resuspended in seawater and injected with dsRNA resuspended in nuclease-free water once a day for three consecutive days. Animals were changed to a fresh dsRNA soaking solution each day following injections (500μL per/well of a 24-well plate, 10-17 worms/well). Animals were either fixed whole, or cut two hours after injection on the last day and allowed to regenerate in fresh seawater for different time points prior to fixation.

### Phospho-histone-H3 (H3P) immunostaining and quantification

H3P staining on fixed worms was done as described in (Srivastava *et al*., 2014). Animals were fixed in 2 mL round-bottom tubes and stored in 100% MeOH at -20°C. Animals were bleached overnight in a 6% H_2_O_2_ solution in MeOH, rocking at room temperature under a strong light source. The following day, animals were washed three times in 100% MeOH, and gradually rehydrated to PBS+0.1% Triton-X-100. Animals were permeabilized in a Proteinase K solution for 15 minutes (0.1% SDS, 20mg/mL Proteinase K in PBS+0.1% Triton-X) and post-fixed in 4% paraformaldehyde in PBS+0.1% Triton-X-100 for 20 minutes. Animals were washed three times for 10 minutes each in a PBS+ 0.1% Triton-X-100 solution. Permeabilized animals were blocked in 10% Horse serum in PBS+0.5% Triton-X-100 solution for one hour at room temperature, and were then incubated overnight at 4°C with primary antibody (Rabbit anti-H3P antibody, Millipore 06-570) at 1:1,000 concentration. Afterward, animals were washed six times for 10-20 minutes each in PBS+0.5% Triton-X-100. Animals were incubated with secondary antibodies at 1:500 concentration (AlexaFluor488 goat anti-rabbit IgG, Jackson Immunoresearch 111-545-144, resuspended 1:1 in 100% glycerol) in block solution overnight at 4°C. Animals were washed six times for 10-20 minutes each in PBS+0.5% Triton-X-100, incubated with DAPI for 45 minutes, mounted using Vectashield as described previously(Srivastava *et al*., 2014), and imaged on a Leica SP8 microscope. Image processing and numbers of H3P^+^ cells were counted using the image processing program Fiji(Schindelin *et al*., 2012). Briefly, confocal image stacks were first z-projected. Images were segmented into four equal sections using the Grid Tool, and the number of H3P^+^ cells within each region was manually counted. Cell numbers were normalized by the area of the fragment, which was measured by drawing a region of interest of each segment, thus providing a measure of the number of mitotic cells per unit area (cells/mm^2^).

### Cell co-expression quantification

Confocal image stacks of stained animals were imaged using a 63X objective, then individual images within each stack were manually assessed for co-expression of *in situ* labels and counted. Images were taken from three representative sections of the animal, across several animals per condition (n > 3). Cells were counted by hand and recorded using the Cell Counter plugin in FIJI.

### Irradiation experiments

Whole animals were exposed to 10,000 rads using a cesium source (our source is 209 rads/minute; animals were exposed to source for 48 minutes in a petri dish filled with seawater). After five days, animals were injected and soaked in dsRNA as described above, then cut and assessed using the *in situ* protocol as described above. Previous studies(Srivastava *et al*., 2014) have demonstrated a loss of all cycling cells following lethal irradiation by 7 days post irradiation.

### ATAC-seq and ChromVAR analysis

ATAC-seq data were collected, analyzed, mapped, and validated as described in(Gehrke *et al*., 2019). ChromVAR analysis was also used as described in(Gehrke *et al*., 2019) to identify EGR binding sites in the *wnt-3* and *wnt-1* genomic loci.

## Supplemental Information

**Supplemental Figure 1.1:**
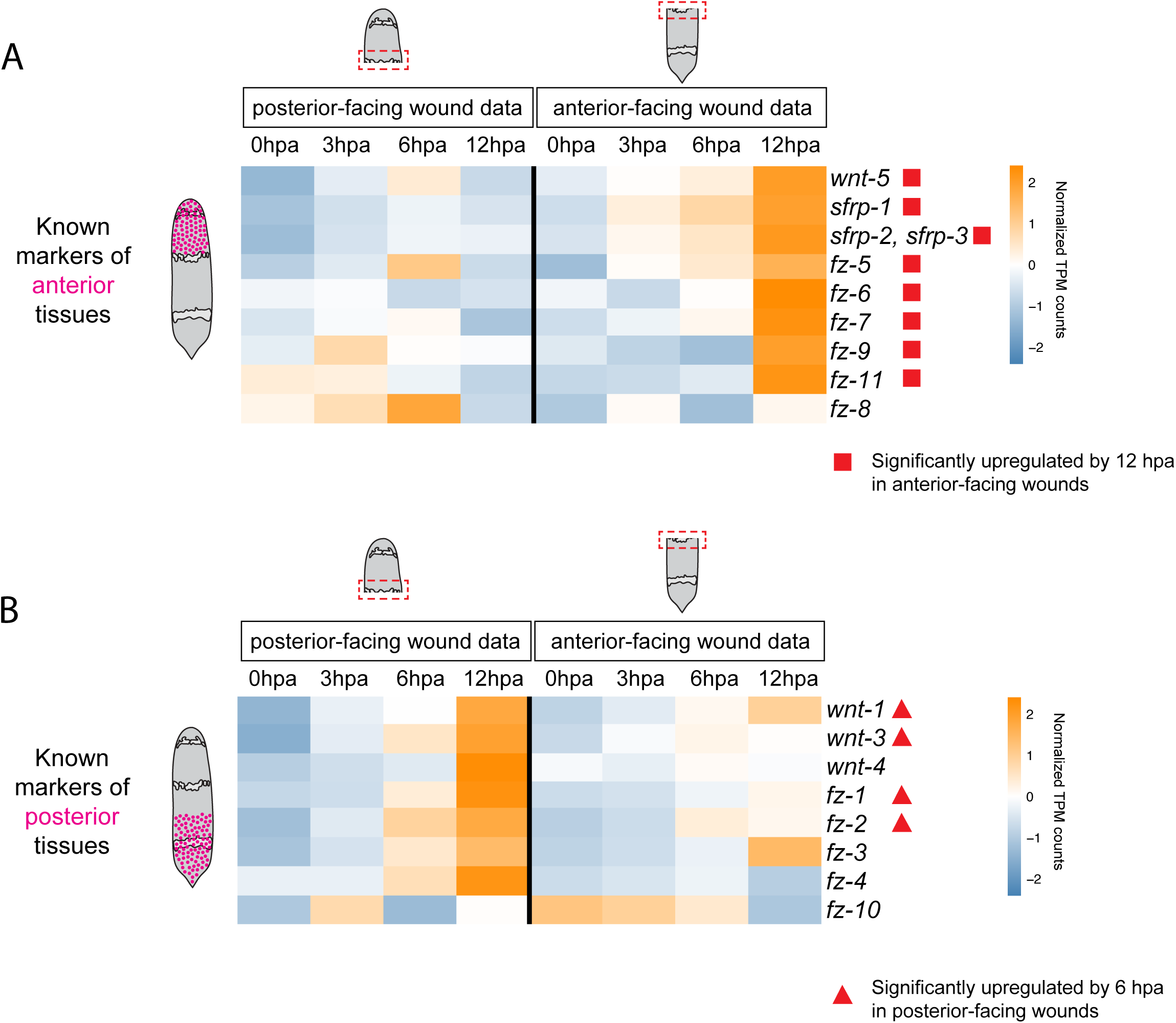
Expression of Wnt ligands during regeneration. (A, B) Expression of known markers of anterior or posterior identity was established by 6-12 hpa in anterior- or posterior-facing wound sites. Heatmap of normalized TPM (transcript per million, compared to 0 hpa) values of published markers with known anterior (A) or posterior (B) *in situ* hybridization patterns in *Hofstenia*. Gene names at right are from (Srivastava et al. 2014).

**Supplemental Figure 1.2:**
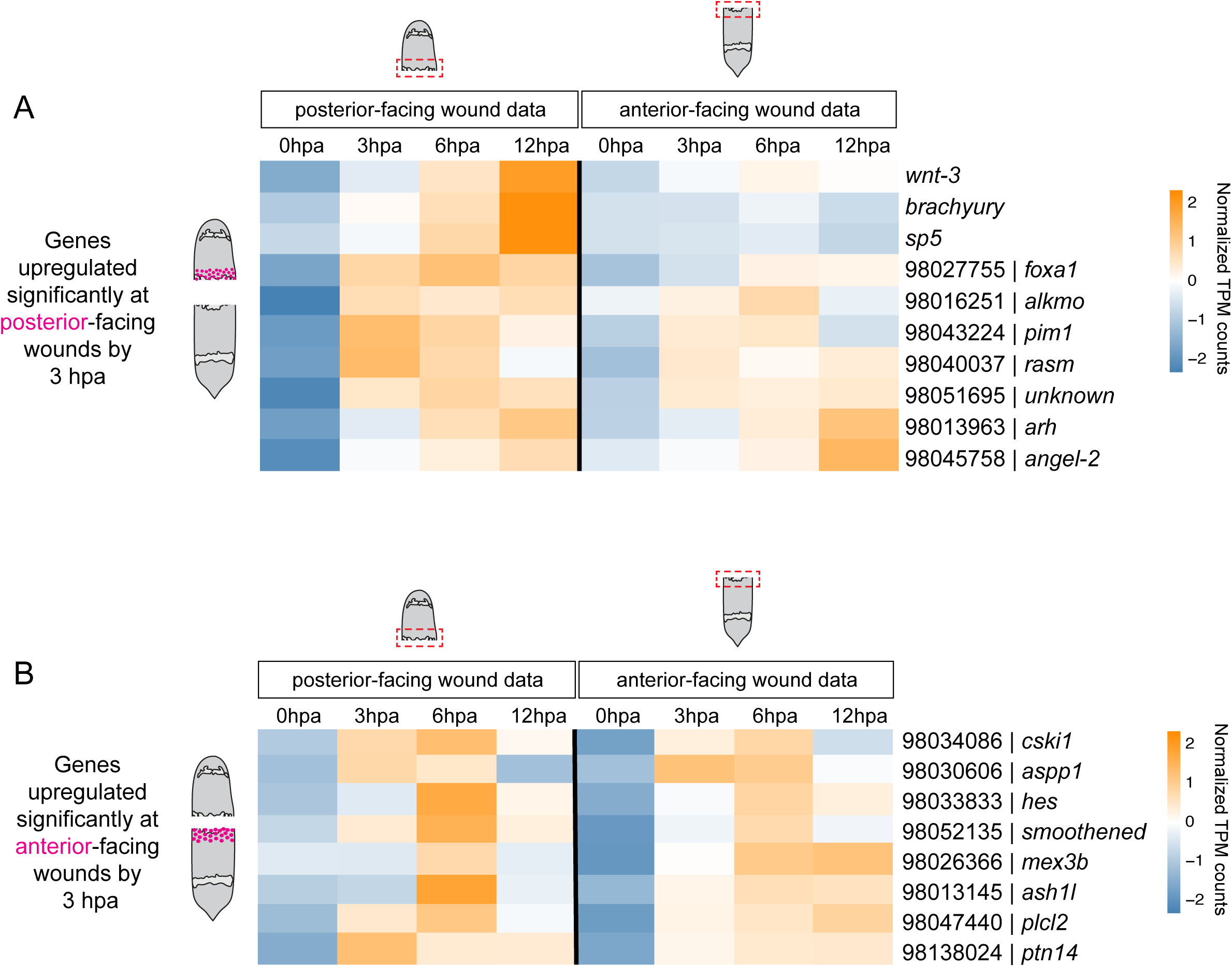
Regeneration transcriptome analysis identified genes with asymmetric expression in anterior- or posterior-facing wound sites by 3 hpa. (A,B) Heatmap of normalized TPM (transcript per million, compared to 0 hpa) values of genes significantly upregulated by 3 hpa in anterior- (A) or posterior-facing (B) wound sites during regeneration identified in our analysis. Normalized TPM values shown from 0 hpa to 12 hpa in both fragments. Gene names depicted were closest human BLAST hit or if previously described, published gene name was used. Datasets from (Gehrke et al. 2019).

**Supplemental Figure 1.3:**
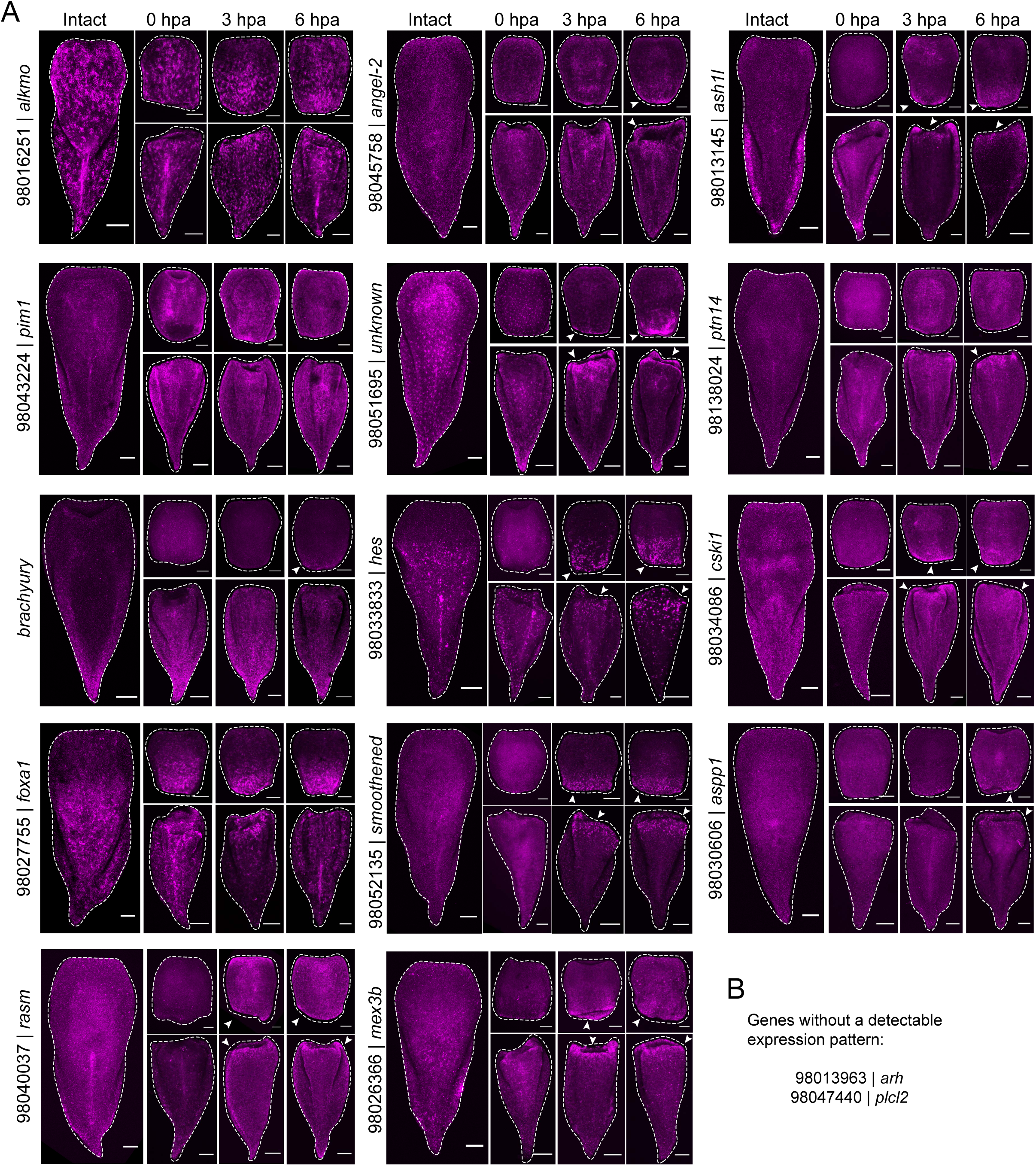
Validation of candidate genes by *in situ* hybridization of regenerating fragments. (A) *in situ* hybridization patterns for all genes we were able to obtain detectable expression patterns for at 0 hpa, 3 hpa, 6 hpa, and in intact animals (16/18). Some genes were expressed at both wound sites (*rasm, angel-2, unknown, smoothened, mex3b, ash1, cski1, aspp1*; white arrowheads) or if only expressed at one wound site, no earlier than 6 hpa (*brachyury, ptn14*; white arrowheads). In a subset of genes, it was difficult to determine if any wound-induced expression was present (*alkmo, pim1, foxa1, hes*). (B) List of the two genes (*arh, plcl2*) we were unable to obtain visible expression patterns for.

**Supplemental Figure 1.4:**
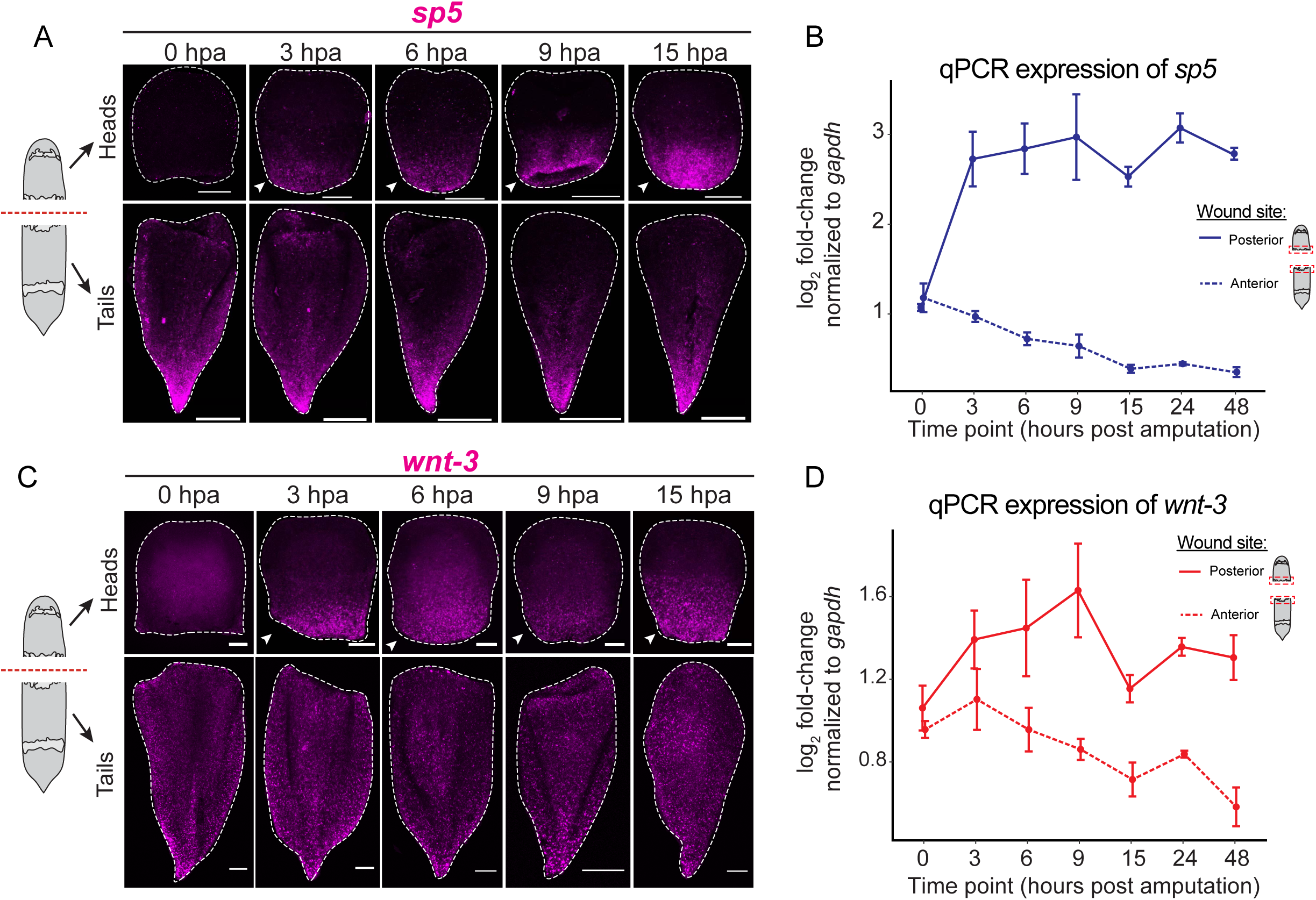
Expression of top candidates (*sp5, wnt-3*) during regeneration by *in situ* hybridization and qPCR. (A, B) Expression of *sp5* in regenerating head and tail fragments by *in situ* hybridization (A) and qPCR (B). *sp5* was expressed in posterior-facing wound sites by 3 hpa, and maintained in this region at later stages of regeneration (arrowheads). (C,D) Expression of *wnt-3* in regenerating head and tail fragments by *in situ* hybridization (C) and qPCR (D). *wnt-3* was also expressed in posterior-facing wound sites by 3 hpa during regeneration, and this expression was maintained at later stages during regeneration (arrowheads). Scale bars, 100μm.

**Supplemental Figure 2.1:**
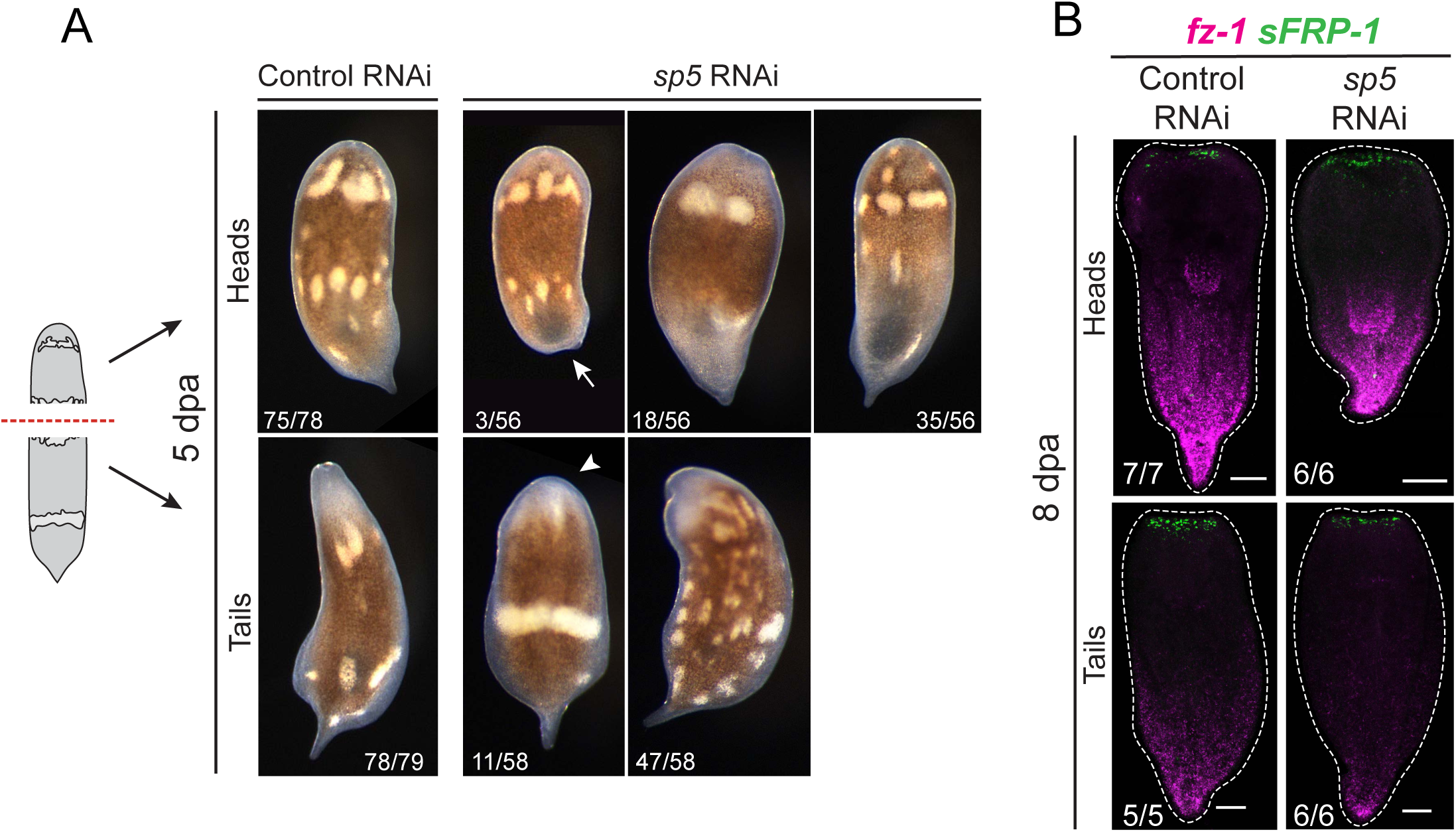
*sp5* RNAi led to a partially penetrant regenerative phenotype. (A) Control RNAi fragments formed a new tail in head fragments (75/78) and an unpigmented blastema in tail fragments (78/79) by 5 dpa. Most *sp5* RNAi head fragments regenerated similar to control RNAi fragments (35/56, heads; 47/58, tails). Some *sp5* RNAi head fragments failed to regenerate a new tail (arrow, 3/56), or formed an outgrowth but not a fully patterned posterior (18/56). Some *sp5* RNAi tail fragments failed to form a mouth or large unpigmented blastema (7/58), or failed to regenerate completely (arrowhead, 4/58). (B) Expression of anterior and posterior markers (*fz-1, sFRP-1*) was unaffected following *sp5* RNAi in both head and tail fragments. Scale bars, 100μm.

**Supplemental Figure 2.2:**
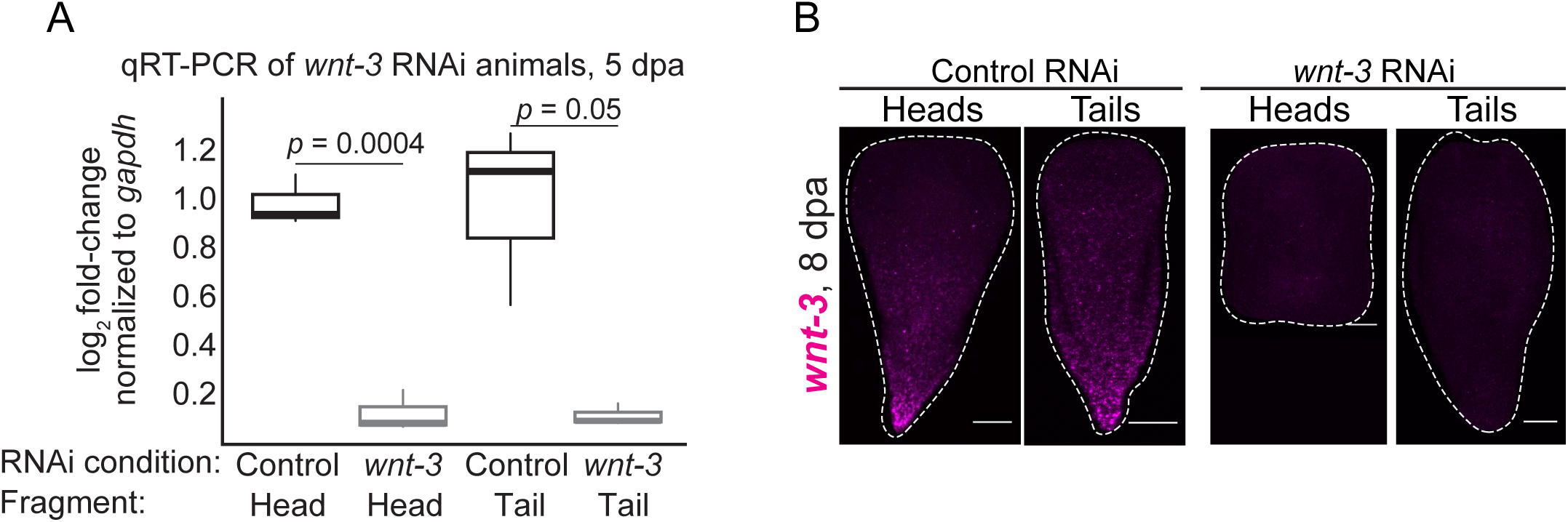
Validation of *wnt-3* RNAi by qPCR and *in situ* hybridization. Expression of *wnt-3* by qPCR (A) and *in situ* hybridization (B) in 5 dpa RNAi fragments. Expression of *wnt-3* was lost following RNAi injections. qPCR samples are n > 3 fragments/replicate, with 3 replicates; *p*-values, Wilcoxon rank-sum test. Scale bars, 100μm.

**Supplemental Figure 2.3:**
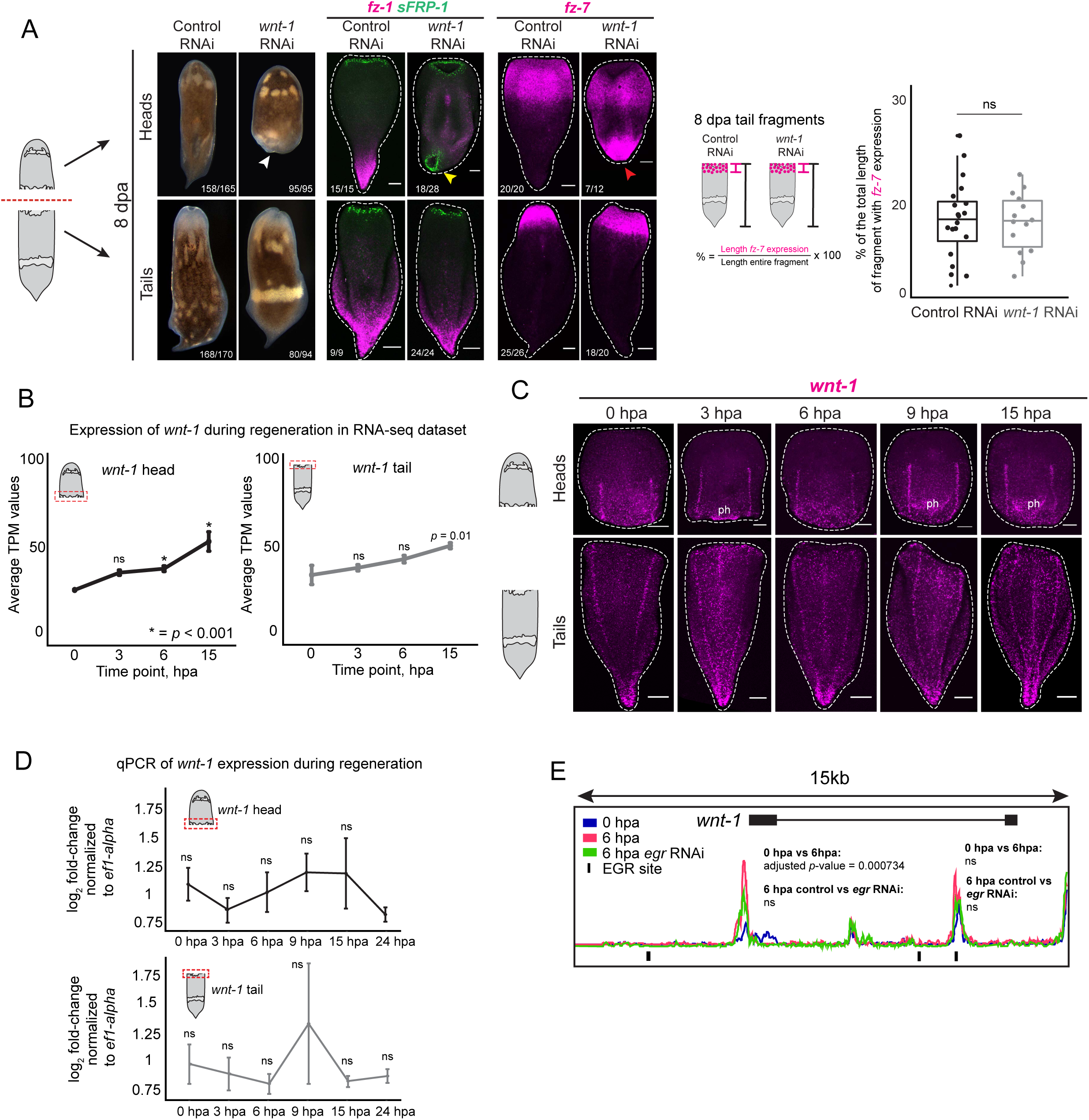
Characterization of the *wnt-1* RNAi regenerative phenotype. (A) RNAi of *wnt-1* caused posterior regeneration defects in head fragments at 8 dpa (white arrow), compared to control RNAi head fragments. *wnt-1* RNAi tail fragments regenerated and formed a visible mouth and blastema in a similar manner to control RNAi tail fragments. Expression of the anterior marker *sFRP-1* in *wnt-1* RNAi head fragments revealed new mouth formation in posterior-facing wound sites (yellow arrow), while anterior-facing wound sites expressed anterior markers in a similar manner to control RNAi. The anterior marker *fz-7* was expressed ectopically at posterior-facing wound sites following *wnt-1* RNAi (red arrow). The expression domain of *fz-7* in tail fragments was the same as in control tail fragments, quantified at left (n > 13 fragments per RNAi condition; *p*-value = 0.85; Welch two-sample *t*-test; ns, not significant). Scale bars, 100μm. (B) Expression of *wnt-1* in the RNA-seq dataset from 0 to 12 hpa. (p-values, likelihood ratio test; * = *p*-value < 0.001). (C) Expression of *wnt-1* by *in situ* hybridization in regenerating fragments from 0 hpa to 15 hpa. There was no visible wound-induced expression pattern of *wnt-1* at any time point during regeneration (ph, pharynx). Scale bars, 100μm. (D) Expression of *wnt-1* by qPCR in regenerating posterior-facing (top) and anterior-facing (bottom) wound sites from 0 hpa to 24 hpa. (E) Genomic locus of *wnt-1*. ATAC-seq data showed a region of chromatin within the promoter of *wnt-1* that became accessible at 6 hpa (magenta) compared to 0 hpa (blue). Accessibility of this region was not sensitive to *egr* RNAi compared to control (green; ns, not significant; Wald test). Control RNAi track omitted for clarity. Accessibility of an additional peak (at right) within this region did not change during regeneration or in response to *egr* RNAi.

**Supplemental Figure 3.1:**
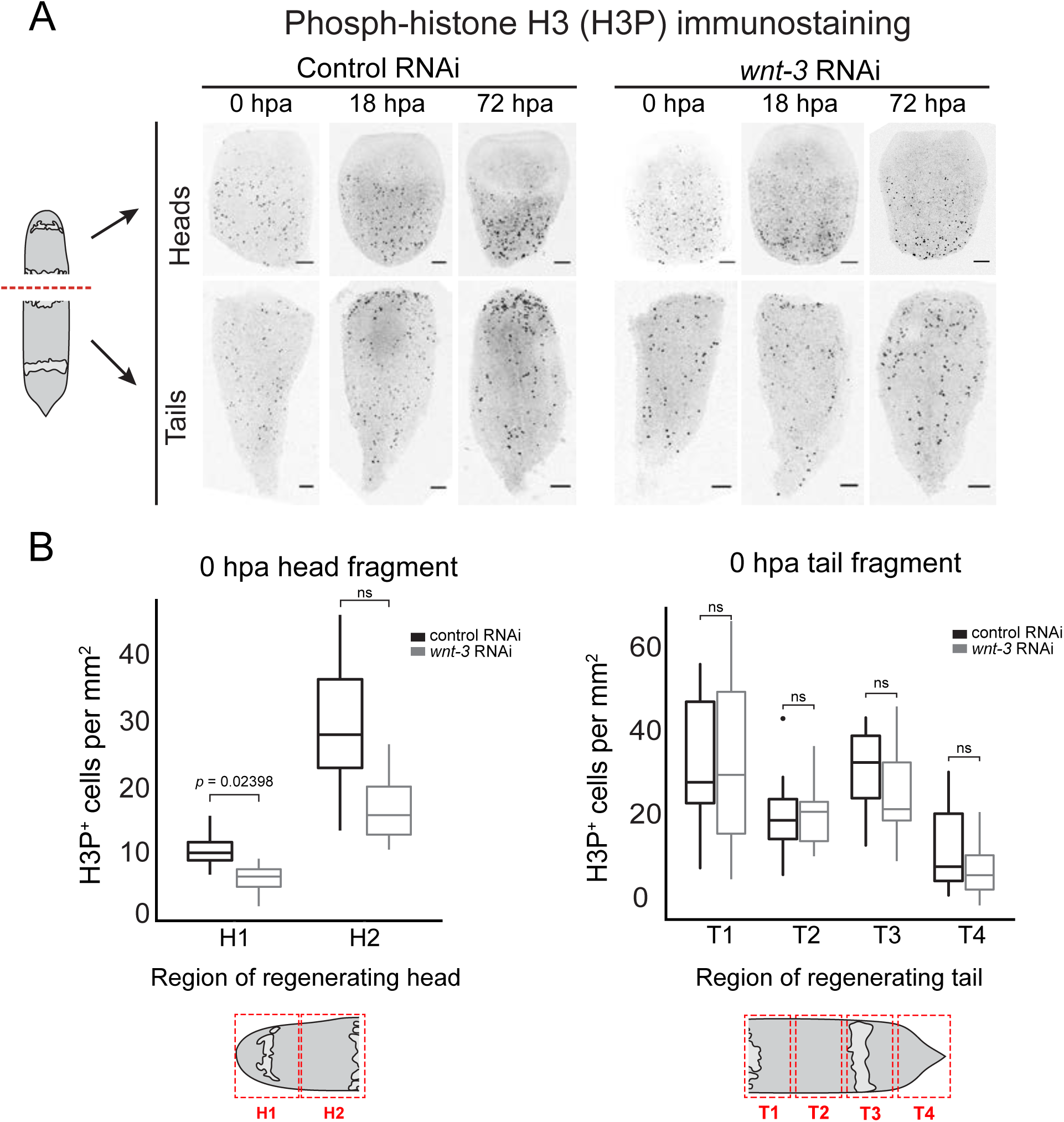
Phospho-histone H3 (H3P) immunostaining and quantification in *wnt-3* RNAi fragments. (A) Representative images of Phospho-histone H3 (H3P) immunostaining in control and *wnt-3* RNAi regenerating fragments at 0 hpa, 18 hpa, and 72 hpa. LUT has been inverted for clarity. Scale bars, 100μm. (B) Quantification of H3P^+^ foci in 0 hpa head and tail fragments following *wnt-3* or control RNAi. (n > 4 per RNAi condition/time point; *p*-value, Wilcoxon rank-sum test; ns, not significant).

**Supplemental Figure 3.2:**
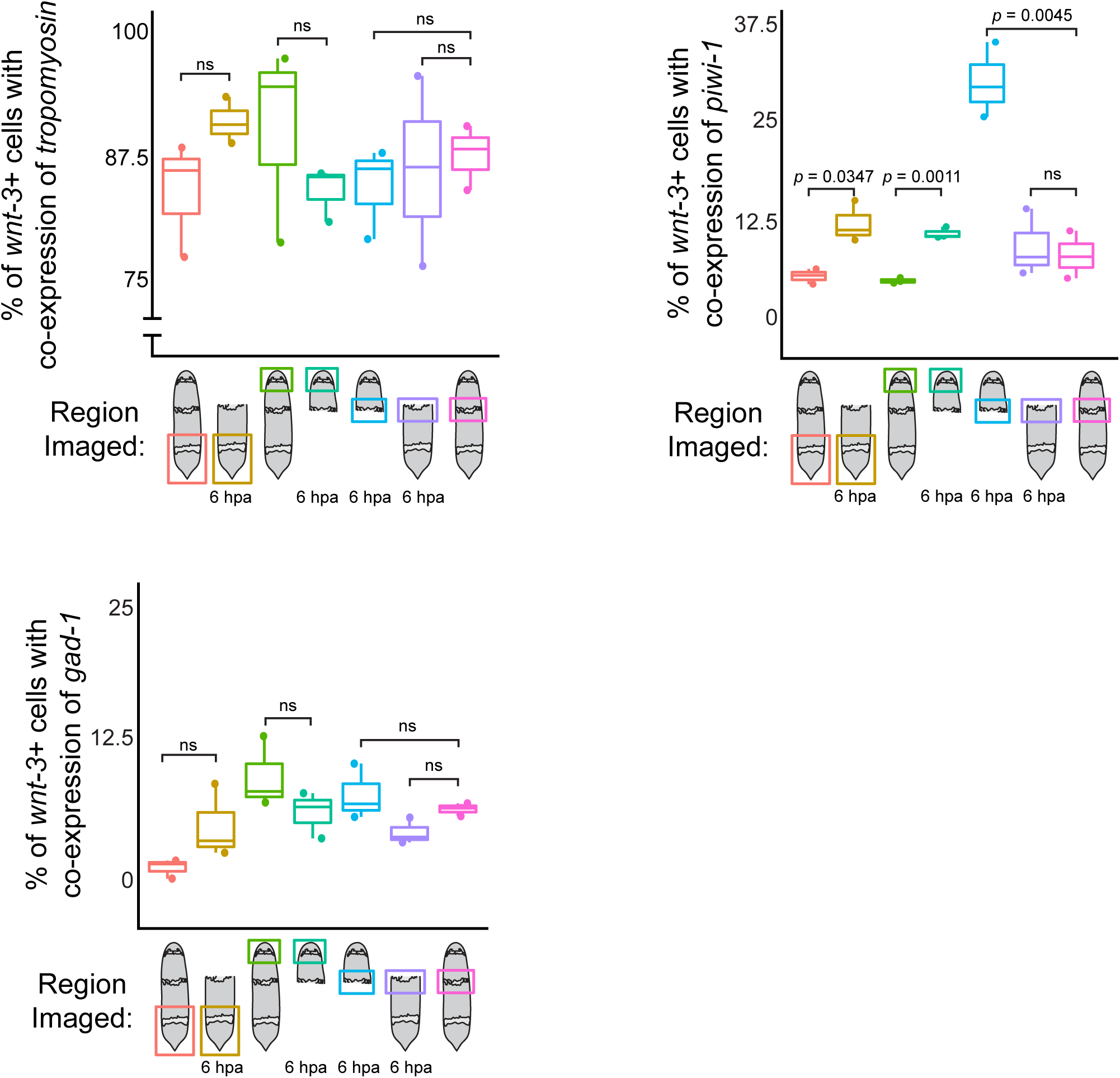
Quantification of co-expression of *wnt-3* with cell type markers. Co-expression was assessed in whole animals and in 6 hpa anterior- or posterior-facing wound sites, and in regions distant from the wound site in 6 hpa fragments. No significant differences were detected in numbers of *wnt-3*^*+*^/*gad-1*^*+*^ cells or *wnt-3*^*+*^/*tropomyosin*^*+*^ cells, regardless of condition or region sampled. Following amputation, co-expression of *wnt-3* and *piwi-1* was significantly increased in 6 hpa anterior- and posterior-facing wound sites compared to equivalent regions of intact animals. (*p*-values, Shapiro-Wilks Normality test, Welch two-sample *t*-test).

**Supplemental Figure 3.3:**
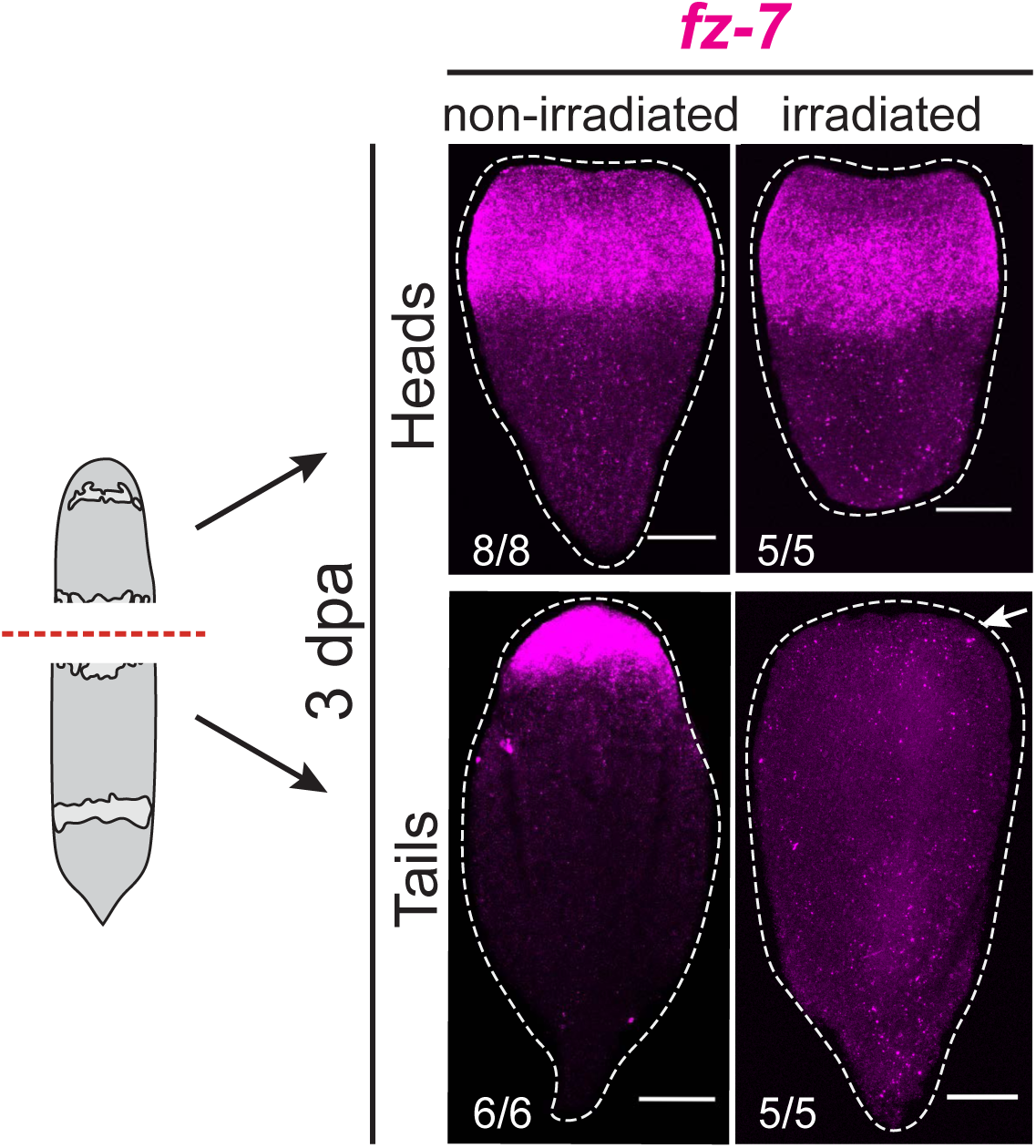
Expression of *fz-7* in wild-type and irradiated regenerating fragments. Wild-type animals expressed *fz-7* in an anterior domain in head fragments and in anterior-facing wound sites of tail fragments at 3 dpa. Tail fragments lost anterior-facing wound sites expression of *fz-7* following irradiation by 3 dpa (white arrow). Numbers of fragments with shown phenotypes in the lower left corner. Scale bars, 100μm.

**Supplemental Figure 5.1:**
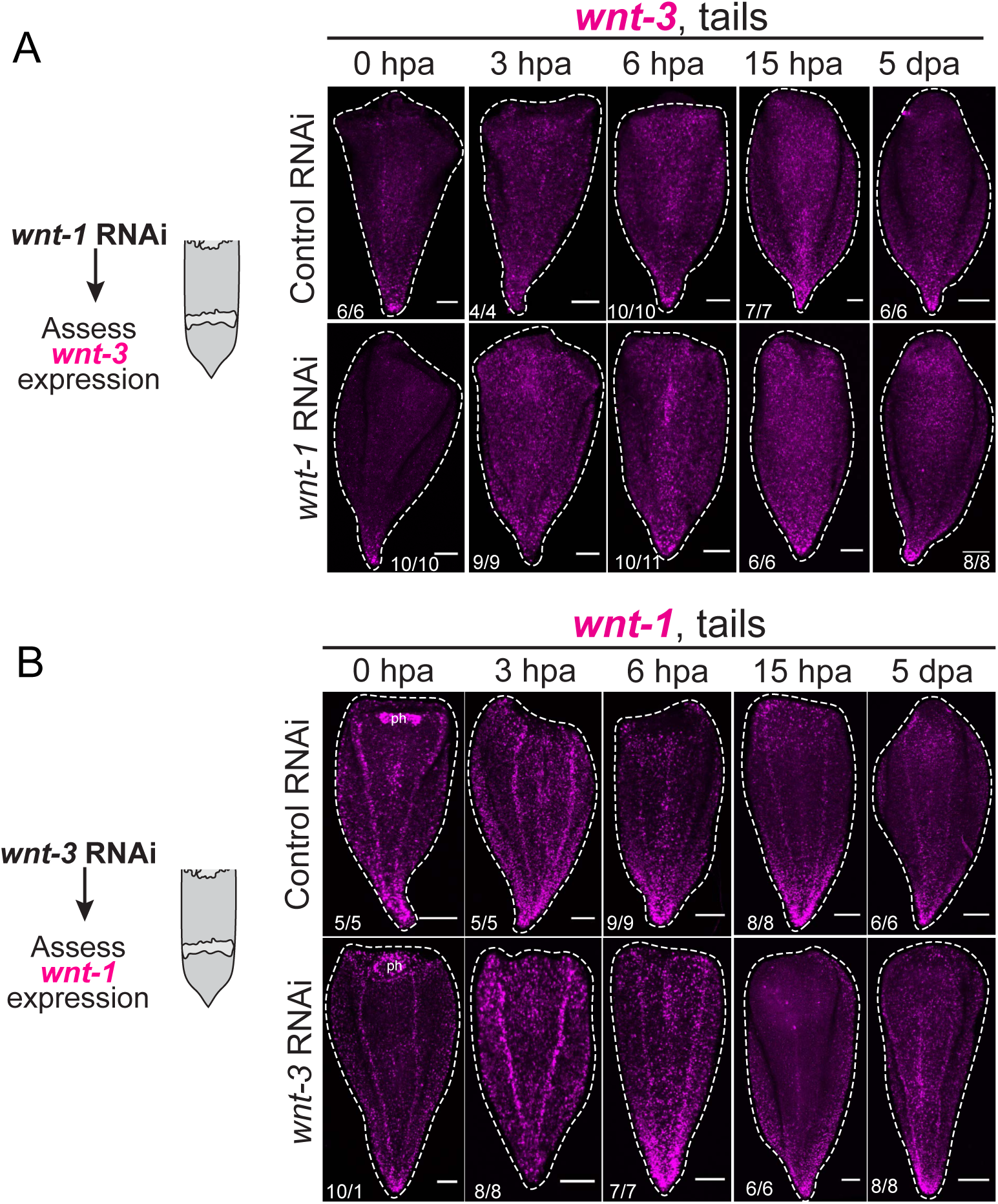
Upregulation of *wnt-3* during regeneration relied on pre-existing expression of Wnt pathway members. (A) Expression of *wnt-3* by *in situ* hybridization in control and *wnt-1* RNAi tail fragments. The posterior gradient of *wnt-3* was maintained in both control and *wnt-1* RNAi tail fragments during regeneration. (B) Expression of *wnt-1* by *in situ* hybridization in control and *wnt-3* RNAi tail fragments. *wnt-1* was expressed in a posterior gradient in both control and *wnt-3* RNAi tail fragments during regeneration (ph, pharynx). Scale bars, 100μm.

**Supplemental Figure 5.2:**
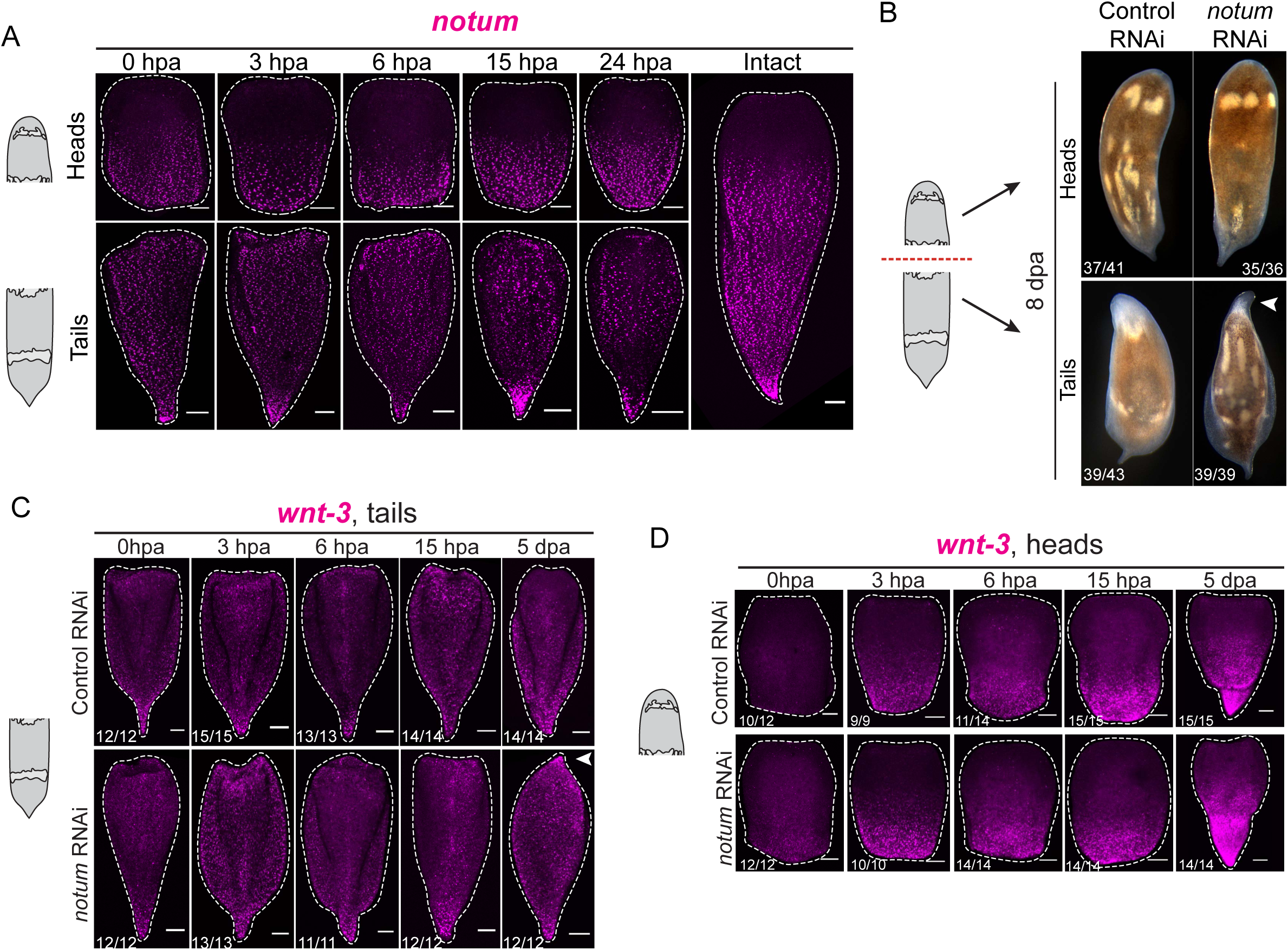
Expression of *notum* during regeneration, RNAi phenotype, and expression of *wnt-3* during regeneration following *notum* RNAi. (A) Expression of *notum* during regeneration by *in situ* hybridization. There was no clear wound-induced expression pattern of *notum* at any time point during regeneration. Scale bars, 100μm. (B) *notum* RNAi head fragments formed a posterior, but tail fragments formed an ectopic tail at anterior-facing wound sites (arrowhead; Srivastava et al, 2014). Numbers of fragments with indicated phenotypes in the lower left corner. (C) Expression of *wnt-3* in regenerating *notum* RNAi tail fragments. Expression of *wnt-3* at early stages of regeneration was unaffected, but by 5 dpa, *wnt-3* was expressed in the ectopic posterior tissues formed at anterior-facing wound sites following *notum* RNAi (arrowhead). Numbers of animals with indicated phenotypes in lower left corner; all scale bars, 100μm. (D) Expression of *wnt-3* in control and *notum* RNAi head fragments. *wnt-3* was expressed in posterior-facing wound sites by 3 hpa in control and *notum* RNAi animals, and this expression was maintained at later stages during regeneration.

## Tables

**Supplemental Table 1:** (A-B) Genes identified in RNA-seq analysis at 6 hpa (A) and 3 hpa (B). (C) Summary of *in situ* hybridization results of all genes assessed. (D) qPCR primers used for validation of expression during regeneration and validation of RNAi knockdown. (E) Raw counts of cells co-expressing tissue markers and *wnt-3*.

